# Cryo-EM structure of gas vesicles for buoyancy-controlled motility

**DOI:** 10.1101/2022.05.08.489936

**Authors:** Stefan T. Huber, Dion Terwiel, Wiel H. Evers, David Maresca, Arjen J. Jakobi

## Abstract

Gas vesicles allow a diverse group of bacteria and archaea to move in the water column by controlling their buoyancy (1). These gas-filled cellular nanocompartments are formed by up to micrometers long protein shells that are permeable only to gas. The molecular basis of their unique properties and mechanism of assembly remains unknown. Here, we solve the 3.2 Å cryo-EM structure of the *B*.*megaterium* gas vesicle shell made from the structural protein GvpA that self-assembles into hollow helical cylinders closed off by cone-shaped tips. Remarkably, the unique fold adopted by GvpA generates a corrugated cylinder surface typically found in force-bearing thin-walled structures. We identified pores in the vesicle wall that enable gas molecules to freely diffuse in and out of the GV shell, while the exceptionally hydrophobic interior surface effectively repels water. Our results show that gas vesicles consist of two helical half-shells connected through a unique arrangement of GvpA monomers, suggesting a mechanism of gas vesicle biogenesis. Comparative structural analysis confirms the evolutionary conservation of gas vesicle assemblies and reveals molecular details of how the secondary structural protein GvpC reinforces the GvpA shell. Our findings provide a structural framework that will further research into the biology of gas vesicles, and enable rational molecular engineering to harness their unique properties for acoustic imaging (2, 3).

## Main

Microorganisms utilise active motility systems to move towards or away from a variety of environmental stimuli such as chemicals and light (4). These include swimming by rotation of rigid flagella; and movement over solid surfaces with filamentous appendages (5). Other forms of motility do not rely on active propulsion. Aquatic bacteria and archaea have evolved mechanisms to regulate buoyancy and can – similar to ballast tanks in submarines – create and eliminate gas-filled compartments to allow vertical migration in the water column. The cellular compartments providing positive buoyancy are formed by gas-filled protein shells called gas vesicles (GVs) (1).

There are very specific requirements for such structures: to achieve net buoyancy, GVs must occupy a substantial proportion of the cell, which involves forming compartments that extend over hundreds of nanometers in size. To maximise buoyancy the shell must be constructed from minimal material. At the same time, the shell needs to provide resistance to hydrostatic pressure to maintain buoyancy with changes in water depth (6). GVs have therefore evolved as rigid, thin-walled structures composed of a single protein unit that typically polymerises into large cylindrical shells closed off by conical tips (7, 8). The shell allows gas to diffuse passively between the GV lumen and the surrounding liquid, while effectively repelling water (9). All GVs identified to date appear to be constructed from the same components (10). The ∼7 kDa primary gas vesicle protein GvpA forms the core of the GV shell and the cone-shaped tips. A second protein, GvpC, binds the exterior of the gas vesicle and provides additional structural reinforcement (11, 12).

With molar masses exceeding hundreds of MDa, GVs range among the largest protein-based macromolecular assemblies reported to date. Despite intensive efforts (7, 13–19), the molecular structure of GVs and therefore a molecular-level understanding of its distinctive properties have remained elusive. Here, we present the native state cryo-EM structure of the canonical gas vesicle shell, providing detailed insight into the biogenesis of GVs and the unique evolutionary adaptions that enable buoyancy-controlled motility.

### Cryo-EM structure of the gas vesicle wall

We expressed and purified *B*.*megaterium* GVs that form narrow tubes most suitable for cryo-EM analysis [Fig. 1, Extended Data Fig. S1a-c]. The native *B*.*megaterium* GV gene cluster contains two almost identical GvpA homologs, named GvpA and GvpB [Extended Data Fig. S2]; for consistency in naming convention, we will refer to them as GvpA1 and GvpA2. A minimal gene cluster with only GvpA2 is sufficient for GV assembly in *E*.*coli* (20, 21) [Fig. 1a]. Cryo-EM images showed GVs forming 0.1-1 µm long cylinders with varying diameters (55±7 nm), consistent with previous data (22) [Fig. 1]. A subset of images (∼16 %) contained GVs with diameters as small as 34-42 nm [Fig. 1b, Extended Data Fig. S3a], which had their cylinder shape best preserved in the thin liquid film of cryo-EM samples [Extended Data Fig. S1a-e]. We used a combination of 2D and 3D classification techniques to select 4% of the small GV subset corresponding to 35.6 nm diameter [Extended Data Fig. S3b] and used helical reconstruction to obtain a cryo-EM density of 3.2 Å resolution [Extended Data Fig. S3c]. The final reconstruction yielded a cylindrical GV shell assembly with ∼93 GvpA2 monomers per helical turn, which represents one member of a range of helical polymorphs with diameters ranging from 34 to 70 nm and with 90 to 183 monomers per helical turn.

**Fig. 1.**
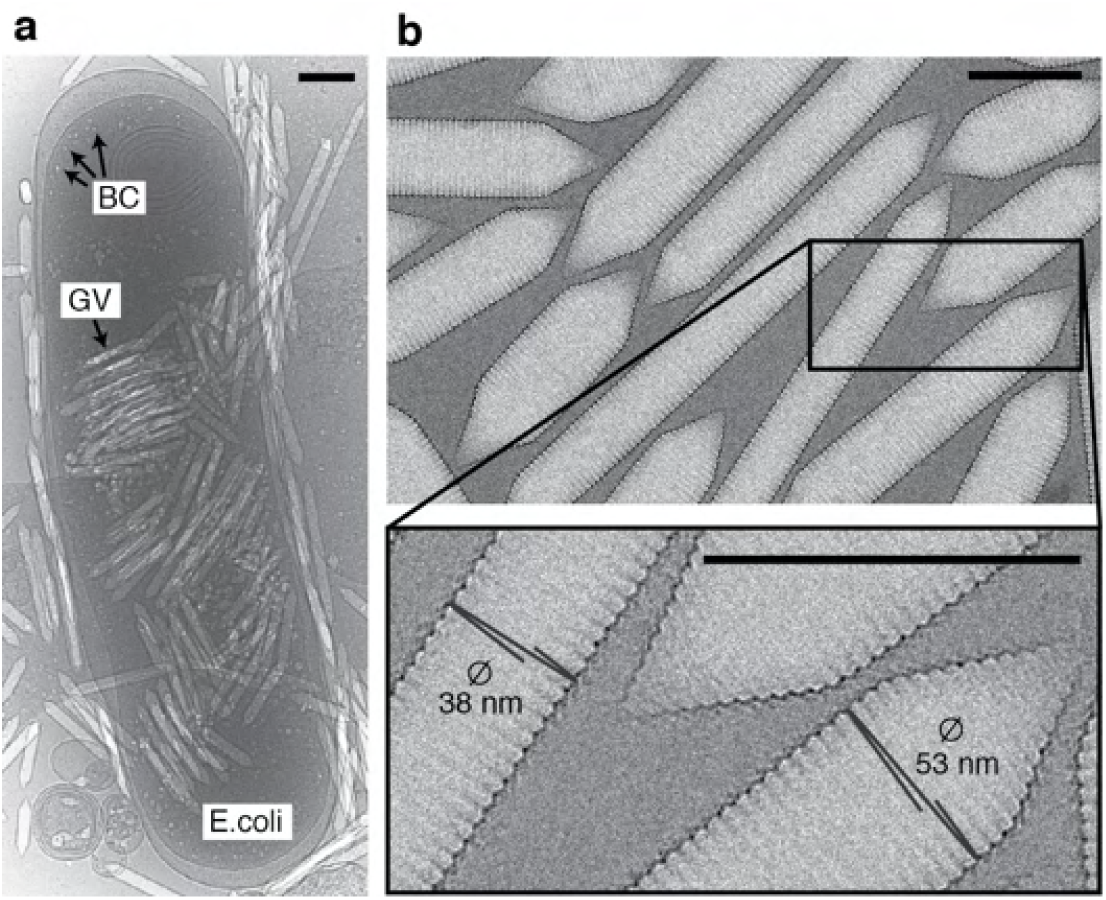
*B*.*megaterium* gas vesicles. (a) Cryo-EM micrograph of an *E*.*coli* cell heterologously producing gas vesicles (GVs). Mature GVs and small bicone (BC) nuclei are visible inside bacteria and in the surrounding medium. (b) Cryo-EM micrographs of purified GVs. GVs appear brighter than the surrounding solvent due to the lower density of the GV-contained gas. The inset shows close-up examples of average-sized GVs (right) and a small diameter GV (left) used for structure determination. All scale bars 100 nm.

The reconstructed density allowed *de novo* atomic model building of GvpA2 in the structural context of its native assembly [Extended Data Fig. S5, Extended Data Table S1]. The cylindrical shell is constructed from thousands of GvpA2 monomers polymerised side-by-side into ribs spiralling into a left-handed helix with a helical pitch of 48.8 Å and -3.87°helical twist, resulting in 92.93 GvpA2 units per helical turn [Fig. 2a,b, Supplementary Movie S1]. Contrary to postulated models (16, 18), GvpA monomers and not antiparallel dimers form the repeating units of the helical assembly. GvpA2 adopts a coil-α-β-β-α-coil fold [Fig. 2a,b]. The carboxyl-terminal residues Asp^67^-Ile^88^ are flexible and not resolved in our structure. The helical lattice forms an array of ribs consisting of densely packed GvpA2 subunits with a lateral centre-to-centre distance of ∼12 Å. The central β-hairpin, tilted at -36°relative to the long axis of the cylinder, forms the core of the GV ribs. Helix α2 folds back onto the hairpin, and helix α1 forms a bridge across the ∼16 Å gap separating adjacent ribs. The GV wall is therefore only one or two peptide layers thick [Fig. 2b]. The inner wall of the GV shell forms a continuous hydrophobic surface consisting of a dense pattern of hydrophobic residues located on the luminal side of the βhairpin and helix α1 [Fig. 2c]. Connections between the ribs of the GV shell are formed by the predominantly polar N-terminus, which extends perpendicular to helix α1 and folds across the βhairpin of the adjacent rib, stabilised by interactions with several residues in the β-hairpin and helix α2 [Fig. 2d, Extended Data Fig. S6].

**Fig. 2.**
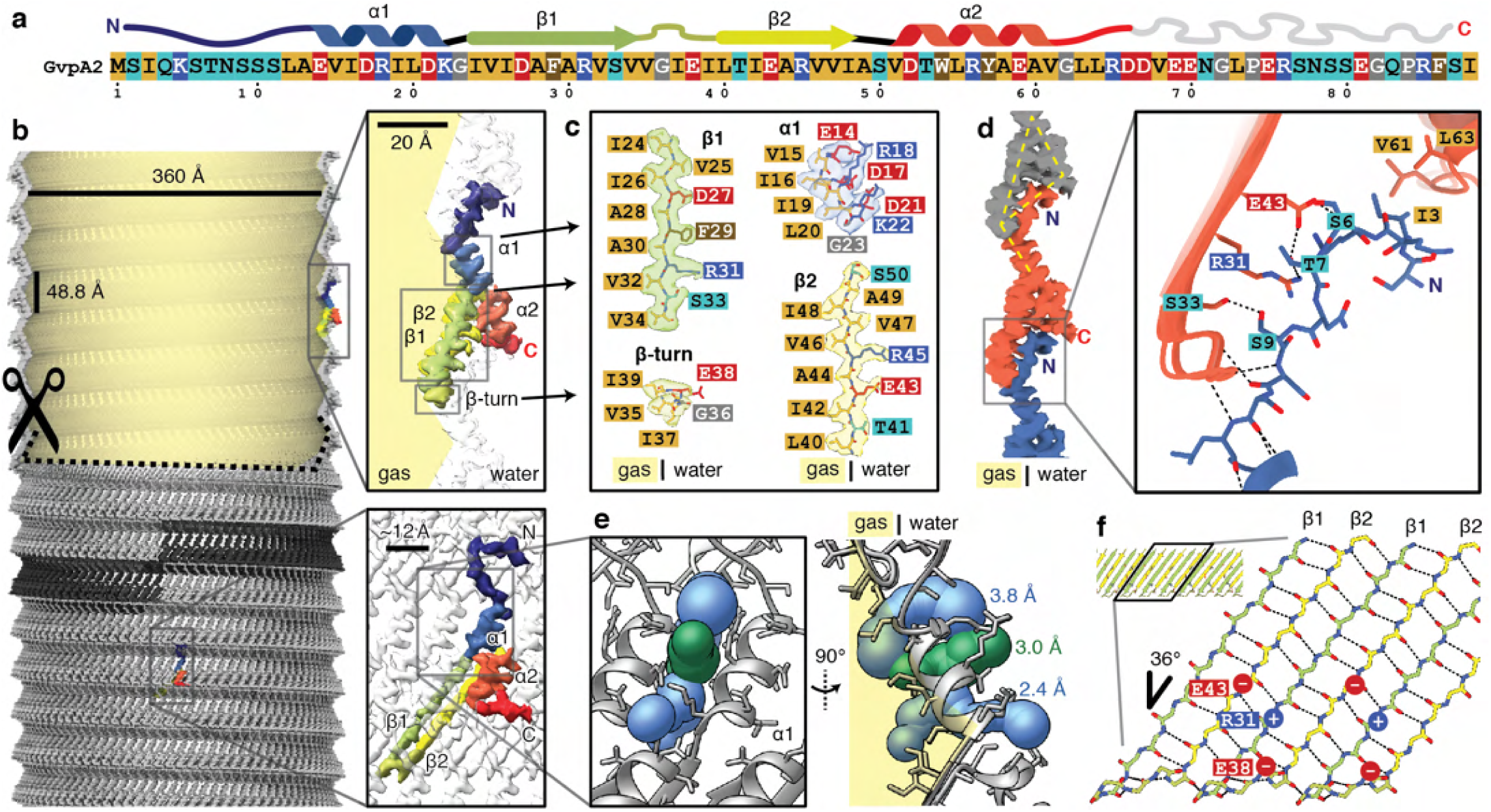
Cryo-EM structure of the gas vesicle wall. (a) Primary and secondary structure of *B*.*megaterium* GvpA2. Residues in the primary structure are coloured based on physicochemical properties. (b) GvpA2 monomers form thin-walled gas-filled protein shells assembled into a left-handed helix. One individual rib formed by 93 monomers is highlighted in dark gray. The top part of the GV is cut open to visualise the gas space. A single monomer is shown in side-view (top inset) and front-view (bottom inset) and coloured based on sequence (N-terminus blue, C-terminus red). (c) Aliphatic residues (Ala,Val,Leu,Ile) line the gas-facing side of the GV wall. (d) Side-view illustrating corrugated zig-zag structure and triangular cross-sections of the wall. Close-up of inter-rib interactions mediated by the N-terminus (blue), which binds across the β-hairpin and the C-terminus (orange) of adjacent ribs, stabilised by hydrogen bonds from backbone and side-chains (Ser^6^, Thr^7^, Ser^9^) and hydrophobic contacts (Ile^3^). (e) A slit between α1 helices allows diffusion of gas through the wall. Three computed tunnels approximate the slit and have bottleneck sizes ranging from 2.4 to 3.8 Å. (f) Schematic of the β-strand rib providing the majority of lateral connections for the assembly through backbone hydrogen bonding (dotted lines) and electrostatic interactions (Glu^43^-Arg^31^-Glu^38^).

### Structural adaptions supporting GV function

The extreme hydrophobicity of the luminal GV surface constitutes an energetic barrier for diffusion of liquid water or condensation of gaseous H_2_O. Consistently, GVs have been shown to be impermeable to water but to be highly permeable to gas molecules (9). How gas molecules passage through the GV wall has so far been unknown. We located pores in the GV shell formed by slit-like openings between α1-helices of adjacent GpvA2 monomers [Fig. 2e, Extended Data Fig. S7]. We quantified the resulting pore size in the GV assembly computationally using Voronoi diagrams (23) and retrieved three different access routes with minimal constrictions ranging from 2.4 - 3.8 Å, compatible with the collisional cross-sections of gases dissolved in the cytosol [Extended Data Fig. S7] (24).

Despite its limited thickness, the GV shell can resist several atmospheres of pressure without collapse (6). GvpA2 monomers are held together tightly by lateral connections along the GV ribs formed by an extensive hydrogen-bonding network between the β-strand backbones [Fig. 2f]. The hydrogen bonds are oriented at an angle of 54°relative to the cylinder axis, which is close to the “magic angle” (54.7°) at which transverse and longitudinal stresses are equal in the wall of a cylinder (25). Additional reinforcements are made by a continuous network of salt bridges formed by Glu^43^–Arg^31^ between two monomers and Arg^31^–Glu^38^ within a monomer. The GvpA2 shell consists of alternating line segments and triangular cross-sections providing force-bearing elements [Fig. 2d]. The corrugations increase stiffness along the rib direction, while the linear segments provide compliant hinge elements that increases elasticity of GVs, increasing their capacity to accommodate deformations orthogonal to the rib without collapse (26).

### Gas vesicles consist of two half-shells with inverted orientation

The thin film of the cryo-EM sample orients the large GVs into a sideways orientation, providing a consistent viewing direction onto the GV edges in projection. Detailed inspection of gas vesicles edges revealed that GvpA2 monomers are always oriented with their β-turns pointing towards the center of the GV cylinder [Fig. 3a], which contains a structural irregularity that has previously been referred to as a seam (15). 2D class averages of GV edges around this seam showed two oppositely oriented GvpA2 monomers that make contact via their β-turns [Fig. 3a] implying that this is the contact sites of two GV half-shells with inverted orientation.

**Fig. 3.**
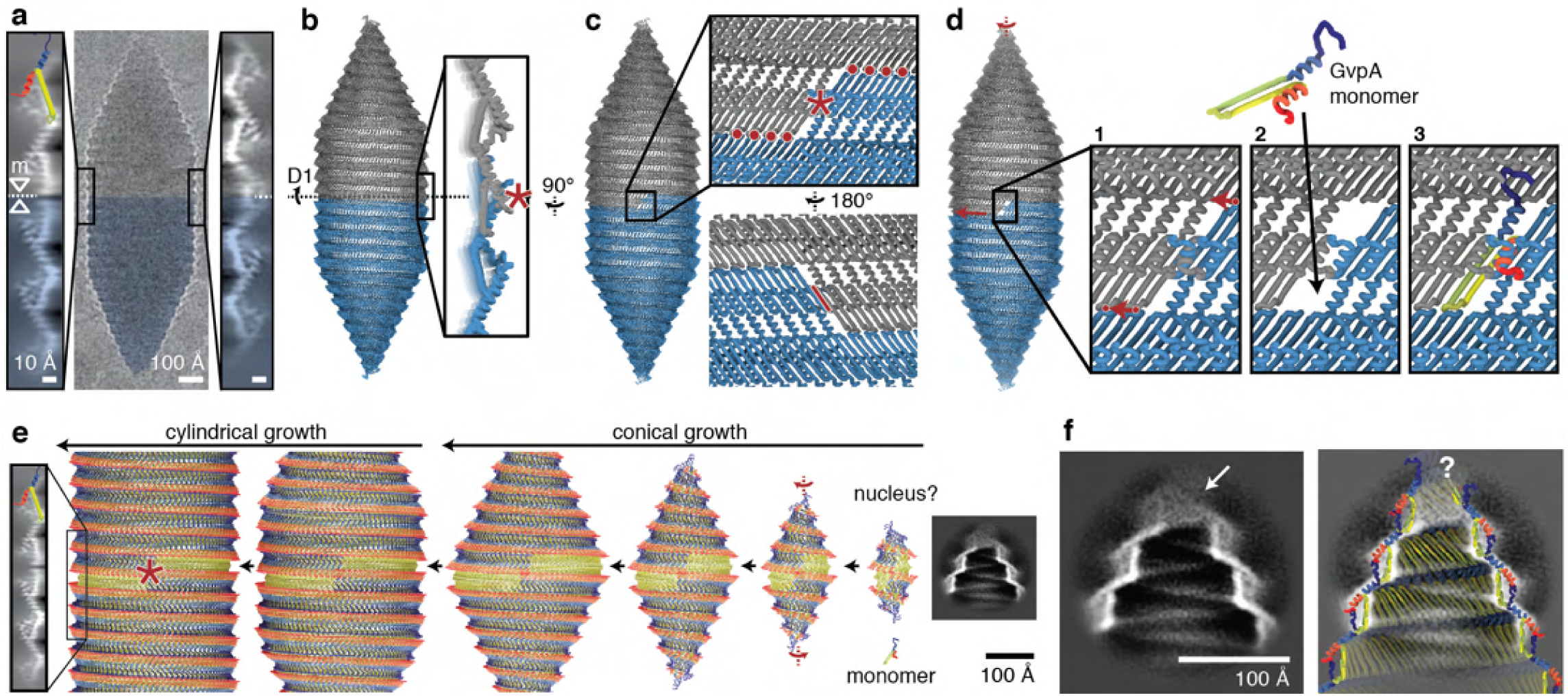
Gas vesicles are assembled from two half shells with inverted orientation. (a) Raw cryo-EM image of a single GV (with inverted contrast). β-hairpins of GvpA2 (cartoon) always point towards the seam at the center of the GV cylinder. Two different types of 2D class averages of the seam (left and right) are observed. The mirror symmetry (mirror axis: m) suggests a 180 °symmetry axis (D1) at the point where two inversely oriented GV half shells meet. (b) Pseudo-atomic model of a GV constructed from two identical halves (gray, blue) with close-up side view of the polarity reversal point (PRP, red asterisk) of the GvpA2 rib. (c) Close-up view onto the Gvp2 lattice with the PRP (red asterisk). Red dots indicate molecular contacts along the GV circumference where β-turns contact. The red line indicates contact between parts of β-strands 2 at the PRP. (d) Model of monomer insertion at the PRP. The two GV halves rotate against each other, with the βhairpin contacts sliding over each other (red arrow) in a ratcheting fashion to allow monomer insertion in the resulting gap. (e) Extrapolation of GV growth to a hypothetical nucleus. Arrows indicate the direction of GV maturation from right to left. A 2D class of GV tips with putative nucleus remnants is shown to scale. (f) Class averages of the GV tips (overlaid with cut-through of the pseudo-atomic model) show no molecular order at diameters lower than 50 Å towards the cone end.

While two contacting cylinders or cones have the same contact geometry along the entire circumference, GVs are assembled from two contacting helices. Continuity of the helicity implies there must be a unique polarity reversal point (PRP) along the circumference, where an upwards and a downwards-oriented GvpA2 monomer meet side-by-side. Apart from classes showing contacting β-turns, there is one less frequent set of 2D classes of the seam that displays two overlapping GvpA2 monomers in inverted orientation [Fig. 3a, Extended Data Fig. S8b]. We posit that this 2D class is a projection view of the PRP located at the GV edge. As GVs can freely rotate around the cylinder axis, the PRP will be located exactly at the edge only at special rotation angles, explaining why this class is observed less frequently. The mirror symmetry in the 2D class implies a viewing direction perpendicular to a 180°(C2/D1) rotation axis [Extended Data Fig. S8c] that points right through the PRP.

Using restraints from our 3D reconstruction and 2D class averages, we constructed a pseudo-atomic model of a GV half-shell. Starting at the PRP, we extendend the model from the D1 symmetry axis using the known helical symmetry for the cylindrical part of the GV shell and allowed transitioning into the conical tips by linearly decreasing the radius set by the 25° semi-angle of the cone and while refining structural adaption of GvpA monomers at defined hinge-points to match the experimental data (details in Methods and Expanded Data Fig. S9). We then duplicated the half-shell by rotating around the D1 axis, leading to a complete GV model consisting of 1730 monomers and a total molecular mass of 12.2 MDa. Simulated density projections from this model closely match the experimental 2D classes [Extended Data Fig. S10a,b]

### Molecular mechanism of gas vesicle biogenesis

Our model of the GV assembly shows that half-shells interact through contacts at the GvpA β-turns around the circumference of the seam, as well as at the PRP where the GvpA2 rib reverses its polarity [Fig. 3c, Extended Data Fig. S10c]. The pattern of side-by-side contacts between β-sheets of GvpA2 monomers (d=d=d=d) along a rib is identical everywhere except at the PRP, where β-hairpins align inversely (d=d-p=p). The PRP is therefore a unique point in the GV assembly that may be recognised by the molecular machinery that facilitates GV growth, and is likely the point at which new GvpA molecules are inserted during GV growth. We propose that insertion of new monomers on either side of the PRP occurs through ratcheting of the two GV half-shells by rotation relative to each other, generating a single monomer gap at the PRP [Fig. 3d]. This would involve breaking the lateral hydrogen bonds between the two monomers around the PRP, along with breaking and re-establishing contacts of the β-hairpins at the seam with no net loss in energy. In our model, two factors suggest that the PRP represents the weakest point in the assembly. First, the two oppositely oriented monomers at the PRP are connected by only 6 hydrogen bonds between segments of strand β2 around Val^47^, as compared to 11 hydrogen bonds formed between monomers along the rib [Extended Data Fig. S10e]. Second, the monomer orientation at the PRP leads to steric hindrance by the α2 helices of the two symmetry-related monomers [Extended Data Fig. S10d]. We propose that the energetic disadvantage of this conformation facilitates opening of the seam for addition of new monomers during growth.

If the GV grows by adding monomers at the PRP, backward extrapolation leads to a transition point at which the seam forms between two conical bases [Fig. 3e]. This is the point where conical growth transitions into cylindrical growth. Extrapolating further leads to a biconical nucleus which must initially form to start GV biogenesis. According to our model, the original nucleus would remain at the cone tips on both half-shells after maturation [Supplementary Movie S2]. The 2D classes and the fitted pseudo-atomic model suggest that N-terminal residues of GvpA2 monomers crowd together at the tip [Fig. 3f]. At diameters lower than 50 Å, the 2D class averages of these cone tips display weak density [Fig. 3f], Extended Data Fig. S11c-d] sealing off the opening. The fuzzy appearance of the density at the tip is indicative of structural heterogeneity in the nucleating monomers.

### Reinforcement of the GV shell by GvpC

Many GV gene clusters contain a second structural gas vesicle protein GvpC, which is absent from our model GV from *B*.*megaterium* (27). GvpC binds on the outside of GVs and increases the critical collapse pressure of GVs (11, 28). While the essential role of GvpC has been firmly established, how it binds to and reinforces GVs has remained elusive. We acquired cryoEM datasets of *Anabaena flos-aquae* GVs and show that they form identical assemblies to *B*.*megaterium* GVs [Extended Data Fig. S12, S13, Extended Note S1]. To investigate the structural role of GvpC, we imaged the GVs in the presence and absence of GvpC and compared 2D class averages of the GV edges [Fig. 4].

**Fig. 4.**
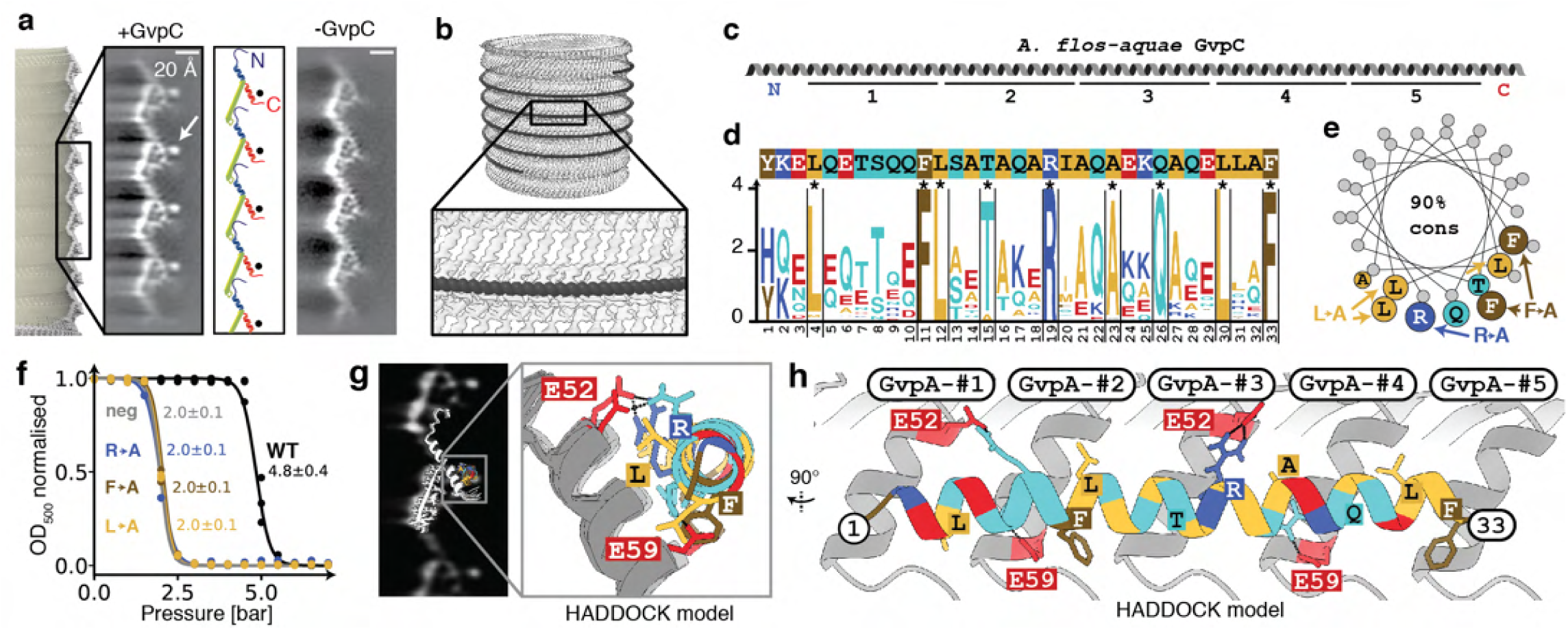
The secondary wall protein GvpC binds along the GvpA ribs of *A*.*flos-aquae* GVs. (a) Comparison of 2D class averages of GV edges with (left) and without GvpC (right) reveal an additional circular density (arrow). A cartoon model of GvpA helps locating the GvpC density to helix α2 (red). (b) Artist impression of GvpC molecules wrapping around GVs. (c) Predicted secondary structure and 33 amino acid repeats 1-5 of *A*.*flos-aquae* GvpC (d) Consensus sequence of the GvpC repeats with logo representation of evolutionary conservation reveals nine highly conserved residues. The height of the characters depict the degree of conservation (information content in bits). (e) Helical wheel plot of highly conserved (>90% conserved in 91 GvpC sequences from other species) amino acids reveal that there is one highly conserved face of the α-helical repeat. Experimentally tested GvpC mutants are indicated. (f) Critical collapse pressure measurements of *A*.*flos-aquae* GVs supplemented with WT GvpC, GvpC mutants or stripped of GvpC. (g) Comparison of 2D class average of GvpC-bound gas vesicles and predicted binding mode between a GvpC consensus repeat and seven *A*.*flos-aquae* GvpA monomers (h) Rotated view of binding model with predicted interactions of residues.

The class average containing GvpC shows an additional dotlike density (∼10 Å diameter) located in vicinity to helix α2 [Fig. 4a]. This feature and its dimensions are consistent with the projection of an α-helix viewed along its helical axis. Indeed, GvpC is predicted to be all α-helical (12) and consists of five 33 residue repeats [Fig. 4b,c] that are highly similar in sequence [Extended Data Fig. S14a]. The class averages suggest that GvpC binds along the ribs of gas vesicles and contacts helix α2 of GvpA [Fig. 4b]. We analysed the evolutionary conservation of the repeat sequence. In a set of 91 GvpC sequences [Extended Data Fig. S14] from different organisms, we find nine residues to be conserved in more than 90% of those sequences, including a strongly conserved set of leucine (Leu^4,12,30^), phenylalanine (Phe^11,33^), glutamine (Glu^26^) and arginine (Arg^19^) residues [Fig. 4d]. A helical wheel plot of the repeat shows that all conserved residues cluster on the same face [Fig. 4e] that likely forms the binding interface. We further confirmed the importance of these residues by designing alanine mutants for occurences in all five repeats [Extended Data Fig. S16, Fig. S17]. All GvpC mutants abrogated stablisation of GVs [Fig. 4f].

Our 2D class averages provide strong spatial restraints on the positioning of GvpC relative to the GvpA ribs, while the conservation pattern and mutants pinpoint residues essential for binding [Extended Data Fig. S14]. We used these data as restraints to predict a model for GvpC binding by computational docking [Fig. 4g, Extended Data S15a-e].

Our data does not allow us to distinguish whether GvpC follows the left-handed spiral of the rib upwards towards the tips, or downwards. We obtained the highest scoring docking solution with the downward orientation [Fig. 4h]. A single GvpC repeat spans four monomers of GvpA. In this GvpA tetrad, glutamate residues Glu^52^ and Glu^59^ in GvpA monomers 1,2, and 4 form hydrogen bonds with GvpC. In our model, Glu^52^ in monomer 3 binds to the conserved Arg19 of GvpC, while Glu^59^ in monomer 4 binds to the conserved Gln^26^. The (Leu^4^, Leu^12^, Leu^30^) triplet inserts between the α2 helices of the GvpA tetrad, while Phe^11^ and Phe^33^ are sandwiched in-between α2 in GvpA monomers 2 and 3 and face-on on helix α2 of monomer 4.

## Discussion

Gas vesicles represent a remarkable example of biomolecular self-assembly. Our results provide a canonical structural framework for the unique molecular properties of gas vesicles, including their selective permeability to gases(9, 29), their mechanical stability (26) and their distinctive ability to grow without compromising the integrity of its shell (1). Our work establishes an atomic resolution model of the mature GV shell formed exclusively by GvpA. A key question is how GvpA nucleates to form an elementary bicone from which the shell extends during GV growth. Besides GvpA, many GV gene clusters contain genes encoding the proteins GvpJ, GvpM or GvpS (27) that exhibit high sequence homology and predicted folds similar to that of GvpA. A dominant structural role of these homologues in mature GVs is unlikely, as none of them were found in intact gas vesicles (30), suggesting a putative role as nucleation factors. Additional support for this role comes from the observation that in some species, GvpJ and GvpM are expressed as part of a separate transcript exclusively during early exponential growth (10), whereas a transcript of gvpACNO is expressed at later stages - possibly to enlarge the formed nuclei. How the initial nucleus is structured and by which mechanism other Gvps assist in growth remain open questions (31).

In most organisms, biconical GVs transition their growth mode when reaching a certain diameter and continue extending cylindrically. This transition occurs over a range of diameters, different for each individual GV. We observed mature cylindrical *B*.*megaterium* GVs with a diameter of 55.5±7.3 nm corresponding to 145±19 monomers per helical turn, and mature *A*.*flos-aquae* GVs with diameter of 87.1±6.9 nm corresponding to 227±18 monomers per helical turn. The mechanism for the bicone-to-cylinder transition is unknown. The GvpA ribs in the cones are highly curved and may be energetically disfavoured. Insertion of new monomers in a bicone results in enlargement of the cone and reduces rib curvature. However, the expanding cone does not provide defined interactions between monomers of adjacent ribs. In contrast, cylindrical GV segments have crystalline order. The interplay of curvature preference and the energetic advantage of crystallinity could favour a cone-tocyclinder transition at a certain critical diameter. This suggests that the GvpA N-terminus, which mediates the rib-to-rib connections, could play a role in determining the size distribution of GVs. Consistently, the sequence of the N-terminus is most divergent. Further evidence for the decisive role of GvpA sequence in defining the final diameter comes from a hybrid GV gene cluster where *A*.*flos-aquae* GvpA integrated into the native gene cluster of *B*.*megaterium*, resulting in GVs with diameters consistent with native *A*.*flos-aquae* GVs (3).

Our structure reveals that the gas permeability of the GV wall can be ascribed to a large number of molecular pores formed between α1-helices of the GvpA shell, the size of which is compatible with the collision diameters (2.65–3.64 Å) of typical atmospheric gases dissolved in the cytosol (24). Surprisingly, the GV wall has been shown to be permeable also to perfluorocyclobutane (C_4_F_8_) with a collision diameter (6.3 Å) exceeding the estimated pore size in our structure. While atmospheric gases appear to diffuse freely through the GV wall (29), the diffusion coefficient of C_4_F_8_ is consistent with a very small number (∼11) of such pores (9). Based on our pseudoatomic model, such pores would need to be located at the PRP or at the conical tips. Alternatively, flexibility in the GV shell may modulate size of the regular gas pores to allow passage of C_4_F_8_.

We show that the five repeats of *A*.*flos-aquae* GvpC bind to α2 helix of the GV wall [see Extended Note S2], thereby increasing the critical collapse pressure. The wall thickness measured between the extremes of the corrugation pattern increases from ∼3 nm to ∼4 nm when GvpC is bound. Because the buckling pressure of thin-walled cylinders is proportional to the third power of the wall thickness (26), the most straightforward explanation for enhanced stability is that GvpC bound on the outside of GVs increases the wall thickness. The real picture is likely more complicated. Our structure reveals that GVs are assembled from two helical half shells that are connected by a seam. Because of its structure, it is conceivable that the seam is a weak-point of the GV where failure will occur first. In this case, GvpC would be most critical to prevent failure at the seam, which questions its requirement elsewhere on the GV shell.

Recently, GVs have been repurposed as genetically encoded acoustic reporters (2, 32). The high contrast in density between gas-filled GVs and surrounding cellular structures makes them amenable to ultrasound imaging (33). While native GVs display little shell deformation under ultrasound exposure, GVs that are stripped of GvpC become less stiff and scatter non-linearily above a certain pressure threshold (34). This behavior enables amplitude modulation imaging and multiplexing of stripped and unstripped GVs in an in vivo context (34, 35). Engineering conditional binding strength of GvpC can transform GVs into biosensors with switchable acoustic properties (36, 37). Our insights into GvpA-GvpC interaction provide a rational basis for designing such sensors. Moreover, the high-resolution structure of the GvpA shell may enable development of designer GVs with custom mechanical shell properties through direct engineering of the GvpA sequence.

Together, our results establish the molecular basis of a widely conserved buoyancy-controlled motility apparatus in aquatic bacteria and archaea. Our study will form the foundation for adressing a multitude of open questions on gas vesicle biogenesis such as nucleation growth, width regulation, function of other gas vesicle gene products in GV assembly and speciesto-species variability of GV gene clusters.

## DATA AVAILABILITY

The refined atomic model of *B*.*megaterium* GvpA2 (GvpB) has been deposited in the Protein Data Bank under accession code 7R1C. The cryo-EM density of a subsection of the *B*.*megaterium* wall (after local refinement) is available in the Electron Microscopy Data Bank under accession code EMD-14238. A symmetrised density of an entire GV cylinder is available under accession code EMD-14340. The presented pseudo-atomic model of a complete *B*.*megaterium* GV is available on Zenodo 10.5281/zenodo.6458345.

## ACKNOWLEDGEMENTS

The pST39-pNL29 plasmid was a gift from Mikhail Shapiro (Addgene plasmid #91696). We thank Cecilia de Agrela Pinto and Tarannum Ara for help with cloning and reconstitution of GvpC constructs. The data on *B*.*megaterium* GVs were collected at the Netherlands Centre for Electron Nanoscopy (NeCEN) made possible through financial support from the Dutch Roadmap Grant NEMI (NWO.GWI.184.034.014). We thank NeCEN operator Willem Noteborn for Nesupport. We acknowledge support by the European Research Council (ERC) under the European Union’s Horizon 2020 research and innovation programme (Grant agreement No. ERC-StG-852880 to AJ), the Dutch Research Council (NWO.STU.018-2.007 to AJ and STU.019.021 to DM), the 4TU Precision Medicine Program, the Medical Delta Ultra HB program and the Kavli Institute of Nanoscience Delft.

## AUTHOR CONTRIBUTIONS

SH, DT, DM and AJ conceived the project. DT and DM provided initial samples; SH and DT purified gas vesicles. SH prepared cryo-EM samples. SH and WE optimised cryo-EM data collection for GVs. SH processed cryo-EM data; SH and AJ analysed cryo-EM data, interpreted the structure, built the atomic model and the mature GV assembly. SH performed bioinformatic analyses and docking experiments; SH and AJ designed GvpC mutants. SH purified GvpC mutants; DT performed collapse experiments. All authors discussed the results. SH and AJ wrote the manuscript; all authors reviewed the original draft and contributed to the final manuscript. AJ supervised contributions by SH and WE; DM supervised contributions by DT.

## COMPETING FINANCIAL INTERESTS

The authors declare no competing interests.

## Methods

### *B*.*megaterium* gas vesicle expression and purification

The purification protocol for Mega GVs was derived from (21). In brief, BL21-DE3-pLysS *E*.*coli* cells were transformed with the pST39-pNL29 plasmid [Addgene 91696] (21) and 1 L of lysogeny broth (LB) containing 0.2% (w/v) glucose was inoculated with 10 mL of overnight culture. The culture was grown at 37°C until OD=0.5 and GV expression was induced with 20 µM IPTG. Following induction, cells were grown at 30°C for 20 hours. The culture was centrifuged at 500 rcf for 2 hours in 50 mL Falcon tubes. The floating fraction was collected with a peristaltic pump. This process was repeated once more. The resulting 25 mL of liquid were lysed chemically by adding 5 ml of SoluLyse reagent per 50 ml of liquid, 0.25 mg/ml lysozyme and 10 µg/ml DNaseI, and slowly rotated for 1 hour at room temperature. GVs were purified in three overnight rounds of floation separation by centrifugation at 800 rcf in 50 mL Falcon tubes. After each centrifugation step, the GV-containing top layer was gently removed with a pipette after which the GVs were resuspended in PBS containing 6M urea (first round), and subsequently in PBS alone. Final concentration was determined as OD_500_=3.1 by optical density measurement at 500 nm against a sonicated blank. The sample was dialysed into imaging buffer (20 mM Tris, 50 mM NaCl, pH=8) prior to cryo-EM sample preparation.

### *A*.*flos-aquae* gas vesicle purification

The purification protocol for gas vesicles from *A. flos-aquae* was derived from (21). Briefly, *A. flos-aquae* (CCAP 1403/13F), also known as *Dolichospermum flos-aquae*, were grown in 250 mL G625 medium complemented by BG11 medium (Sigma C3061) for approximately 2 weeks until confluence. The culture was centrifuged at 350 rcf for 4 hours or until a floating layer of cells was observed. Subnatant was removed using a syringe before lysing the cells in 500 mM sorbitol and 10% v/v lysis buffer (SoluLyse) at 4°C for 6-8 hours while gently rotating. GVs were purified by three rounds of flotation separation with 4-6 hour centrifugation at 350 rcf. After each centrifugation subnatant was removed by syringe after which GVs were resuspended in PBS at pH 7.4.

### GvpC mutant purification

Codon-optimised genes for *A*.*flosaquae* wild-type and mutant (R19A;F-11,33-A; L-4,12,30-A) GvpC (Uniprot: P09413) including a C-terminal ‘GSGSGS’ linker and a C-terminal 6xHis-tag in a pET-28a(+) vector were obtained from Genscript (New Jersey, United States). The mutations were engineered in all five repeats. Proteins were expressed in *E*.*coli* BL21-DE3 cells grown in 750 mL autoinduction medium (38) for 3 h at 37°C before the temperature was lowered to 20 °C for additional 20 h. The bacteria were harvested by centrifugation and lysed by freeze-thaw cycles in lysis buffer (20 mM Tris, 500 mM NaCl, 0.1% Triton-X, 20 mM imidazole; 5 mL per gram of pellet). Lysozyme (0.15 mg/mL) and DNAseI (10 µg/ml) were added and the lysate rotated for 2 h at room temperature. Isolation of inclusion body and IMAC purification was performed as described previously (21). Protein purity was assessed by SDS-PAGE and concentration was determined according to Bradford.

### Preparation of GVs with recombinant GvpC and collapse pressure measurements

*A. flos-aquae* GVs were stripped of GvpC by resuspending in 6 M urea and 60 mM Tris buffer, using 3 rounds of flotation separation as previously described (21). Recombinant GvpC was added to stripped GVs according to the formulation: 2 x OD_500_ x 198 nM x GV volume (liters) = nmol GvpC. GvpC will be present in a twofold excess under the assumption of a 1:25 molar ratio of GvpC/GvpA (21). The urea solution was then slowly replaced with PBS at pH 7.4 by 2 rounds of 12 hours of dialysis over a 7-10 kDA MWCO dialysis membrane. Finally, 3 rounds floatation separation at 350 rcf removed traces of urea. GVs were diluted to OD_500_ 0.1-0.4 for collapse pressure measurements. Samples were loaded in a pressure vessel and hydrostatic pressure was increased in increments of 0.5 bar using pressurized nitrogen gas. Samples were allowed to equilibrate for 5 seconds after pressure changes before absorption was measured using a spectrophotometer (Ocean optics) at 500nm. OD_500_ values were normalised between the minimum and maximum for each measurement. Three independent re-additions per GvpC mutant were performed and measured.

A sigmoid function with *p*_0_ as the inflection point and *k* as the width was fitted to the curves using the means and standard deviations of measured triplicates (n=3) as input for the scipy ‘curve_fit’ function.

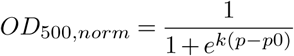

The error of *p*_0_ was determined using a bootstrapping approach, performing the fit 50 times with a random set of n out of n measurement points with replacement. The parameters and their uncertainty were estimated as their mean and standard deviation over the 50 fits.

### Cryo-EM of *B*.*megaterium* gas vesicles

*B*.*megaterium* gas vesicles at OD_500_=3.1 were applied to a freshly glow-discharged Quantifoil R2/1 grid and frozen using a Leica plunger set to 95% humidity, front-side blotting and 20°C with blot times ranging from 5-11 seconds. Micrographs were collected on Titan Krios (Thermo Fisher Scientific) microscope at the Netherlands Center for Electron Nanoscopy (NeCEN) operated at 300 kV. Dose-fractionated movies were acquired on a Gatan K3 direct electron detector at a pixel size of 1.37 Å with 60 frames over an exposure of 30 e^-^/Å^2^ and a defocus range from -0.25 to -1.25 µm.

### Cryo-EM of *A*.*flos-aquae* gas vesicles

Native *A*.*flos-aquae* gas vesicles (containing GvpC) at OD_500_=13 in imaging buffer (20 mM Tris, 50 mM NaCl, 0.5 mM EDTA, pH=8) were applied to a freshly glow-discharged Quantifoil R2/1 grid and frozen using a Leica plunger set to backside-blotting, 95% humidity and 20°C with 10 s blot time. 1273 cryo-EM micrographs at 1.288 Å pixel size were acquired on a JEOL 3200 microscope with a K2 detector using 62 etotal exposure over 60 frames. *A*.*flosaquae* GVs stripped from GvpC (OD_500_=1, in PBS buffer) were prepared and imaged in a similar same way, using 17.6 eover 50 frames, 1.288 pixel size, acquiring 58 micrographs.

### Data processing and structure determination of *B*.*megaterium* GVs

4351 collected super-resolution movies were 2x binned and motion-corrected in RELION 3.1 (39). CTF determination was perfomed using Gctf 1.06 (40). 709 micrographs containing thin GVs with a diameter of 42 nm or less were identified manually. 1021 tubes were manually picked in RELION by selecting start and end coordinates. 36295 overlapping segments with 512 pixels were extracted along the cylindrical sections with a step size of 49 Å (2x binned). 2D classification was done in RELION3.1 with the ‘ignore CTFs until first peak’ option turned on. The resulting 2D class averages were grouped by projecting them along the helical axis and calculating the rim-to-rim distance [Figure S3b] between the two density maxima. A class with 35.6 nm edgeto-edge distance was selected containing 2911 segments. Analysis of in-plane rotated power spectra of the segments using Helixplorer (http://rico.ibs.fr/helixplorer/) revealed a likely helical symmetry between 90.92 to 95.92 units per helical rung, with 49 Å helical pitch. 2D classification was not sufficient to separate all symmetries and the final set of segments originated from several assemblies with different symmetries. 3D classification starting from a featureless cylinder and imposing candidate symmetry parameters was used to further select for segments adhering to a particular symmetry, leading to 1460 segments with symmetry 92.93 units per turn. 3D refinement of those particles with CTF refinement and Bayesian polishing led to a resolution of 3.6 Å at FSC=0.143. Convergence of 3D refinements was only achieved when using a ‘tau_fudge’ parameter of 5..

A final round of particle polishing was used to create a new particle stack extracted only from frames 1-20 (0-10 e^-^/Å^2^) of the movies. Particles were exported to cryoSPARC 3.1.0 (41) and high-pass filtered to 100 Å. 3D refinement with the helical reconstruction algorithm implemented in cryoSPARC also led to a reconstruction at 3.6 Å. To account for small deviations from helical symmetry, e.g. by flattening of the tube in ice, a mask encompassing 3×9 GvpA2 monomers was created in ChimeraX (42). The particle stack expanded by helical symmetry was subjected to focussed refinement in cryoSPARC using the mask, which increased the final resolution to 3.2 Å at FSC=0.143. The final maps were cropped from a box size of 512^3^ voxels to a box size of 128^3^ voxels centred on the refined region.

### Atomic model building and refinement

To build an atomic model of a GvpA2 monomer the final map density was traced de novo using COOT (43). The monomer was expanded using helical symmetry (rise: 0.525 Å, twist: -3.87°) into a 15 subunit segment (three ribs with 5 monomers each) to account for connections between monomers and manually adjusted in ISOLDE (44) before automatic real-sapce refinement in PHENIX 1.13 (45) using NCS restraints between monomers. Renderings of the cryo-EM density and atomic models were made in ChimeraX 1.4 (42).

### 2D classification of edges, seams and tips

From the *B*.*megaterium* gas vesicles cryo-EM dataset, several hundred particles of either seams or tips were picked manually and used to generate a template for automated picking in cryoSPARC 3.3 (41). Particles were high-pass filtered to 100 Å to eliminate the large negative contrast of the gas space in the GV interior. The picked particles were cleaned up by several rounds of 2D classification to give a clean set of particles of either the seam, the polarity reversal point or the tips. These particle sets were used to train a neural network for particle picking (TOPAZ v0.23, (46)), which was then applied to the micrographs to pick seams, PRPs, and tips. Those particles were cleaned by several rounds of 2D classification and led to the final presented 2D classes. For display, the 2D classes were treated in ImageJ using an ‘unsharp mask’ filter.

The two cryo-EM datasets of *A*.*flos-aquae* gas vesicles with and without GvpC were used to obtain 2D class images of the edges leading to side-views of the wall. Movies were imported into cryoSPARC v3.3 (41), motion-corrected and CTF-estimated. For both dataset, frames were used only until an exposure of ∼15 e^-^/Å^2^ because shrinking of GVs was observed for high exposures leading to GV edges blurring out. A few hundred edges were manually selected for 2D classification to generate picking references for the cryoSPARC filament tracer. Particles were extracted with 192 pixel box size and high-pass filtered to 100 Å. Several rounds of 2D classification and selection of sharp classes led to the final 2D classes of the GV edges with and without GvpC. For display, the 2D classes were treated in ImageJ using an ‘unsharp mask’ filter. Similarly, 2D classes of the GvpA lattice from collapsed GVs were calculated from the *A*.*flos-aquae* GV dataset above including GvpC. For this, the dataset was preprocessed in RELION 3.1 (39), points on the lattice manually picked to create a 2D class, which was then used for automated particle picking. Particles were extracted with a box size of 128 pixels and aligned with 2D classification to generate a view onto the collapsed GV wall. The biggest class containing ∼34,000 particles was selected and was treated in ImageJ using an ‘unsharp mask’ filter.

### Pore analysis

Gas pores in the gas vesicle wall were analysed using MOLE 2.5 (23). The gap between α1 helices has a slit-type shape, enabling multiple possible routes for gas diffusion. Several start and endpoints of tunnels around both sides of the slit were selected and tunnels were computed with MOLE. The slit was modelled by three tunnels and displayed in Chimera (47). The minimal constriction of the tunnel was calculated as the diameter of the smallest sphere in the respective tunnel model.

### Bioinformatics analysis

The sequences of the five 33 AA repeats of *A*.*flos-aquae* GvpC (Uniprot P09413) was converted into a consensus sequence ‘YKELQETSQQFLSA-TAQARIAQAEKQAQELLAF’ using the software ‘cons’ from the EMBOSS package (48). The sequence was used as input for the ConSurf Server (49) to search the UniRef90 database with the HMMER web server (50) using one iteration, resulting in 91 sequences. The resulting sequence alignment was displayed and consensus sequences calculated in MView (51). A sequence logo was calculated from the sequence alignment using WebLogo (52). Helical wheel plots were generated in Heliquest (53).

### HADDOCK modelling of GvpC binding to *A*.*flos-aquae* gas vesicles

A α-helical model of the 33 residue consensus repeat of *A*.*flos-aquae* GvpC (‘YKELQETSQQFLSATAQARI-AQAEKQAQELLAF’) was generated in ChimeraX (42) using ideal α-helical backbone dihedral angles ϕ=-57° and Ψ=-47°. A homology model of the *A*.*flos-aquae* GvpA monomer was generated using SWISS-MODEL (54) based on the structure of *B*.*megaterium* structure GvpA2 and the *A*.*flos-aquae* sequence for GvpA (UniProt: P10397). The homology model was extended with the symmetry parameters from the *B*.*megaterium* assembly in ChimeraX (42) into a rib of 7 adjacent monomers.

Both models were used as input for HADDOCK 2.4 (55). For GvpC, residues 4,11,12,15,19,23,26,30,33, which all are >90% conserved in the bioinformatics analysis were chosen as active residues. For GvpA, residues 51-61 of GvpA, part of helix α2 and adjacent to the GvpC density in the 2D classes, were chosen as active residues. All remaining settings were used at default values.

The highest scoring cluster of the docking solutions (HAD-DOCK score: -108.4 +/-5.0) showed GvpC binding across several GvpA monomers along the helical spiral of the rib, with the GvpC sequence oriented inversely to the direction of helical propagation (with the direction from seam to tip). A very similar cluster was found shifted by one GvpA monomer laterally with the same molecular contacts and was discarded. A second, lower scoring type of cluster (HADDOCK score: -90.7 +/-4.6) showed GvpC binding mode with the GvpC sequence oriented aligned to the direction of helical propagation. Docking clusters were displayed in ChimeraX (42) and hydrogen bonds between GvpA and GvpC highlighted with the ‘hbonds’ command. The highest-scoring solution was manually fitted into a 2D class of the *A. flos-aquae* GV wall with GvpC.

### Pseudo-atomic modelling of a complete GV

The model was generated from the solved atomic model of *B*.*megaterium* GvpA2. The GvpA2 monomer was placed next to the x-axis (D1 axis) manually such that a 180°symmetry copy operation around the x-axis would reproduce a side view of the GV edge compatible with the determined 2D class average.

A left-handed parametric helix in 3D space was defined with the parameter *t* corresponding to turns of the helix:

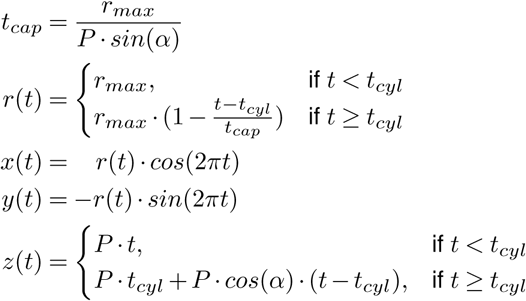

where *t*_*caps*_ is the number of turns in the cap, *t*_*cyl*_ is a userparameter of how many cylindrical turns the model should have (5), *P* is the pitch of the helix of 48.8 Å, *r*_*max*_ the radius of the assembly of 178.4 Å, and *α* is the cone angle of the tip (25°).

The starting point of the curve was adjusted to go through the pivot point (in the center of the two β-sheets between amino acid (AA) 28 and AA 42 carbonyl oxygen atom) of the placed monomer by applying a shift along the z-axis and rotation around the z-axis. Points were placed along the curve at a distance of 12.07 Å, calculated as 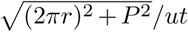, where *ut* is the number of monomers per turn defined by the solved helical symmetry (92.93). 835 points were placed with the last 4 points towards the tip being omitted.

A model only with placement of monomers does not reflect the experimental 2D classes well. Four additional parameters were introduced rotating and modifying the monomer:

- a correction for the change of helix angle towards the ends of the tip as the helix becomes steeper with narrower radius. The rotation center is between the AA28 and AA42 carbonyl oxygen atoms, and the rotation axis normal to a fitted plane through C-αatoms of all amino acids of the βsheets and hairpin (AA23-49).
- a rotation of the entire monomer to account for the tilt of monomers following the cone angle. The rotation center is carbonyl oxygen 36 and the axis normal to the plane of AA24-33 C-βatoms.
- a hinging motion of the two βsheets to account for the deformation visible in 2D classes of the polarity reversal point where GvpA ribs from both gas vesicles halves meet. The rotation axis is defined by a line between the AA23 C-αatom and the AA49 C-αatom, and moves the atoms of AA23-49.
- a hinging motion of the α-helix 1 and the N-terminus, to account for gaps in the assembly formed when monomers tilt towards the cone angle. The axis is defined as in (3), and the rotation moves the atoms of AA2-23.

For the purpose of illustrating GV growth from nuclei to mature cylinders, the model was replicated in Blender 3.0 in a simplified form (without modifications of the monomer) and animated using varying parameters for the diameter and the *t*_*cyl*_ parameter.

## Extended Data

### Supplementary Notes

Note 1 Cryo-EM of *A*.*flos-aquae* GVs and conservation of GV shell architecture. Note 2 Binding of five consecutive GvpC repeats.

### Supplementary Movies

SM1 Movie_SM1_Wall.mp4 - Structural protein GvpA2 in the context of the gas vesicle wall.

SM2 Movie_SM2_Growth.mkv - Artist impression of GV growth from a small nucleus to a mature cylindrical GV.

## Supplementary Note 1: Cryo-EM of *A*.*flos-aquae* GVs and conservation of GV shell architecture

We employed cryo-EM to image native GVs directly purified from cultures of the cyanobacteria *Anabaena flos-aquae* [Extended Data Fig. S12a]. GVs formed cone-ended cylindrical GVs with mean diameter of 87±7 nm, consistent with previous observations (22). 2D class averages of GV edges showed a corrugated zig-zag pattern of the ∼2 nm thick wall formed by GvpA. Another set of 2D class averages obtained from collapsed GVs also present in the images revealed the GV wall to consist of a periodic array of ribs consisting of dense 5.0 × 1.25 nm GvpA subunits tilted at -36°relative to the long axis of the cylinder. The detailed secondary structure visible in the classes showed that individual ribs are formed by β-strands aligning side-by-side. α-helical densities bridge the ∼16 Å gap separating adjacent ribs [Extended Data Fig. S12b]. A cumulative Fourier spectrum computed from in-plane rotated GV segments showed a typical helical transform [Extended Data Fig. S12c], confirming predictions (8) and consistent with our 3D reconstruction from *B*.*megaterium* GVs [Fig. 2]. We therefore attempted to solve the 3D structure of the cylindrical GV wall using helical reconstruction. Using the Fourier spectra together with GV diameter and estimates from the *B*.*megaterium* reconstruction we inferred a range of likely helical parameters by correlation of simulated and experimental Fourier spectra [Extended Data Fig. S12d]. Despite significant effort, 3D refinements of the structure using helical reconstruction did not converge. Closer inspection of full-length GVs revealed them to flatten in the thin ice of the cryo-EM sample, breaking symmetry assumptions of helical reconstruction [Extended Data Fig. S1a]. Nevertheless, the GvpA lattice obtained from 2D classed of longitudinal views (GV edges) and saggital views (2D crystalline GvpA monolayers from collapsed GVs) are indistinguishable from equivalent projections of the *B*.*megaterium* GV structure, hence confirming the conserved architecture of GvpA assembly of the GV shell [Extended Data Fig. S13c,d]. Our results also establish that 2D classification of cryo-EM data can give valuable structural insight into gas vesicle architecture for cases when 3D reconstruction is out of reach. This is due to the small unit cell of the GV wall, where not many features overlap in side views at the GV edges. 2D classes can be calculated to sufficient resolutions to see secondary structure elements and even large side chains. We suggest that this approach can be used for comparative studies of GVs from different species with larger sequence divergence to reveal different assembly modes, e.g. resulting from binding modes of the GvpA N-terminus.

## Supplementary Note 2: Binding of five consecutive GvpC repeats

We inferred a binding mode of a single 33 amino acid repeat of GvpC to *A*.*flos-aquae* GV shells. What remains open is how the five consecutive GvpC repeats are structured. It is likely that all repeats would form identical molecular contacts with the GvpA ribs. The side-by-side distance of GvpA monomers measured between identical points of the inner β-hairpins is 12.1 Å. Due to the curvature of GVs, this distance increases radially outwards across the GV wall towards the binding site for GvpC. For an average 87 nm *A*.*flos-aquae* GV this distance is ∼12.7 Å. A single perfectly α-helical repeat of GvpC would span 49.5 (33 × 1.5Å/residue), only slightly shorter than the 50.8 Å spanned by a stretch of four GvpA monomers. The five tandem repeats of native GvpC would bind to 20 GvpA monomers. Taking into account additional space for the N and C-termini of GvpC, this corresponds well to a previously determined 1:25 molar ratio of GvpA to GvpC (12). The modest distance mismatch between one GvpC repeat and a GvpA tetrad could be accomodated by slight deviation of GvpC from perfect helicity. It has been previously suggested that this would also be required to maintain the relative orientation of consecutive repeats, as with perfect helicity they would be (100°x 33) % 360 = 60°rotated towards each other (12). A small unstructured stretch would allow GvpC to adapt to different curvatures of the GvpA ribs in cylindrical and conical parts and in GVs of different diameters.

**Table S1.** Cryo-EM data collection and model refinement statistics

|  |  |
| --- | --- |
| **Data collection** |  |
| Microscope | Titan Krios |
| Magnification | 64,000 |
| Voltage (kv) | 300 |
| Electron exposure (e<sup>-</sup>/Å<sup>2</sup>) | 30 |
| Exposure time (s) | 2.4 |
| Number of fractions | 60 |
| Number of movies | 4351 |
| Defocus range (µm) | 0.25 - 1.25 |
| Pixel size (Å) | 1.37 |
| Detector | K3 |
| Dose rate detector (e<sup>-</sup>/pix/s) | 24 |
| **Data processing** |  |
| Helical rise (Å) | 0.525 |
| Helical twist (°) | -3.874 |
| Final no. of asym. units | 135,780 |
| Global map resolution (Å)<br>(FSC = 0.143) | 3.2 |
| Map sharpening B-factor (Å<sup>2</sup>) | 60.8 |
| **Model refinement** |  |
| Model starting point | *de-novo* |
| Sequence | GvpB - Uniprot O68677 |
| Model composition |  |
| Non-hydrogen atoms | 497 |
| Protein residues | 65 |
| Validation |  |
| MolProbity score | 0.69 |
| Clashscore | 0 |
| Rotamer outliers (%) | 0 |
| Ramachandran plot |  |
| Favored (residues) | 61 |
| Allowed (residues) | 1 |
| Disallowed (residues) | 1 (V35) |

**Fig. S1.**
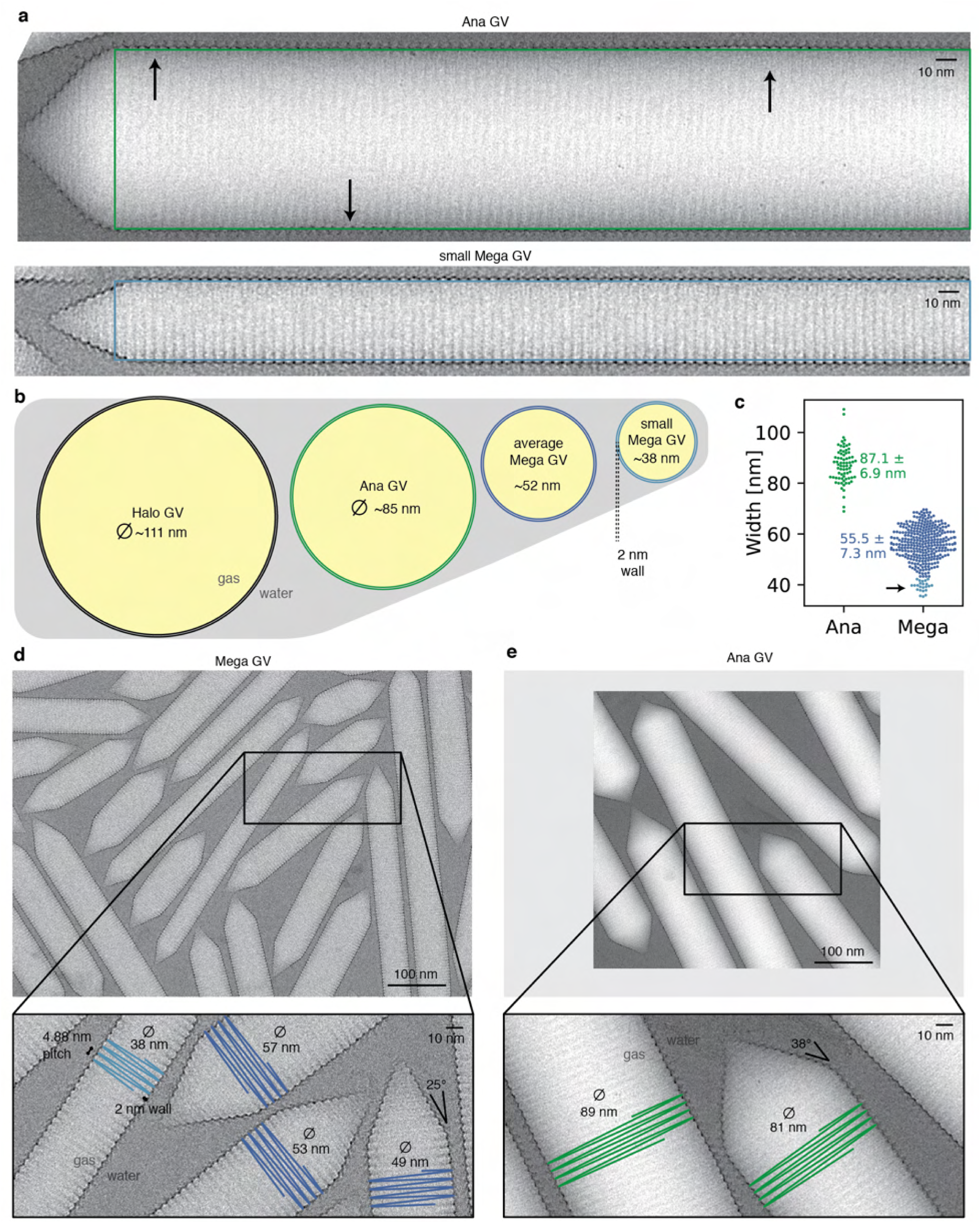
Gas vesicles from different organisms have different mean diameters. (a) *A*.*flos-aquae* GVs with mean diameter of 85 nm (22) deform in cryo-EM sample preparation and do not stay perfectly cylindrical. The green rectangle depicts the ideal, non-deformed shape. Arrows point at deviations of GV from the ideal shape. Helical reconstruction in cryo-EM assumes perfect helical crystals of the repeating unit. This assumption is violated when GVs ‘squish’ during sample preparation. Very thin *B*.*megaterium* GVs maintain a cylindrical shape in the thin ice layer of the cryo-EM sample. (b) Schematic cross-sections of *H*.*salinarum, A*.*flos-aquae* and *B*.*megaterium* gas vesicles drawn to scale. Average diameters from (22). (c) Width measurements in 20 micrographs from both *A*.*flos-aquae* and *B*.*megaterium* GV datasets are in close agreement with those diameters. A long tail of small-diameter outliers with 34-42 nm diameter was observed (black arrow) in *B*.*megaterium* GVs. (d) Representative cryo-EM micrograph of Mega GVs with inset indicating the diameter of several GVs, the 4.88 Å helical pitch, the 2 nm thick wall and the 25°cone angle. (e) Representative cryo-EM micrograph of Ana GVs with inset indicating diameters and the 38°cone angle.

**Fig. S2.**
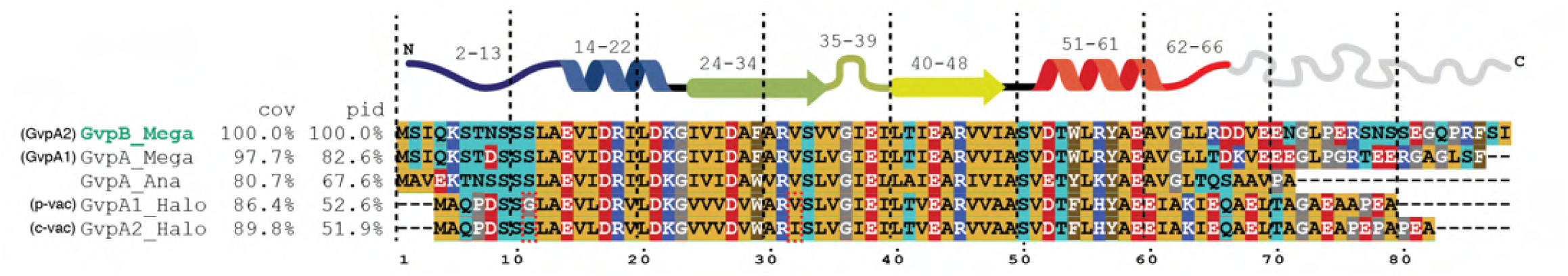
Similarity of the GV wall protein GvpA from selected organisms. Sequence alignment of Mega GvpA (GvpA1) and GvpB (GvpA2), Ana GvpA and Halo GvpA1 and GvpA2 show high degree of conservation despite forming gas vesicles of different diameters.

**Fig. S3.**
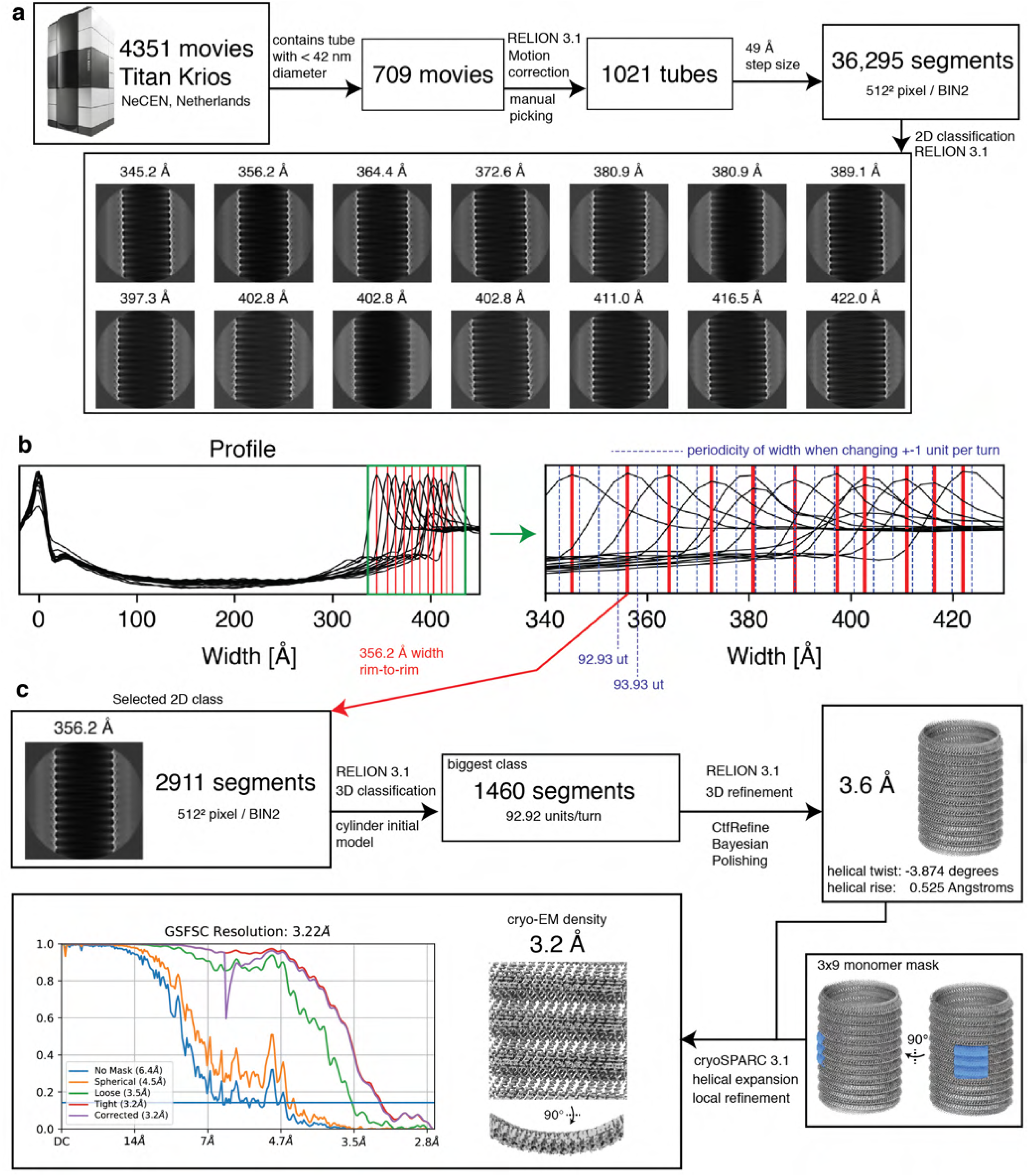
Processing of *B*.*megaterium* gas vesicle dataset. (a) Preprocessing, manual picking, segment extraction and 2D classification leads to 2D class averages of gas vesicles with different diameters. (b) The 2D classes were projected along the helical axis to generate profiles. The profiles were aligned with respect to the left peak. Zooming into the right peaks shows the distribution of gas vesicle widths in the 2D classes. Peaks are marked with a vertical red line. Blue lines indicate the periodicity of widths when an increment of one monomer per helical turn is assumed, based on a side-to-side distance of monomers of 12 Å, leading to a diameter increment of 12/π=3.8 Å. Two to three different helical polymorphs are part of the particle subset belonging to a single 2D class average. (c) Processing steps starting from 2911 selected segments of a particular 2D class. The particle subset from the 2D class was further selected by 3D classification, imposing possible symmetry candidates between 90 and 95 units per wrung to select 1460 segments. Focussed refinement on a 3×9 monomer segment of the wall in cryoSPARC 3.1 (41) leads to the final result of a 3.2 Å resolution cryo-EM density of the gas vesicle wall.

**Fig. S4.**
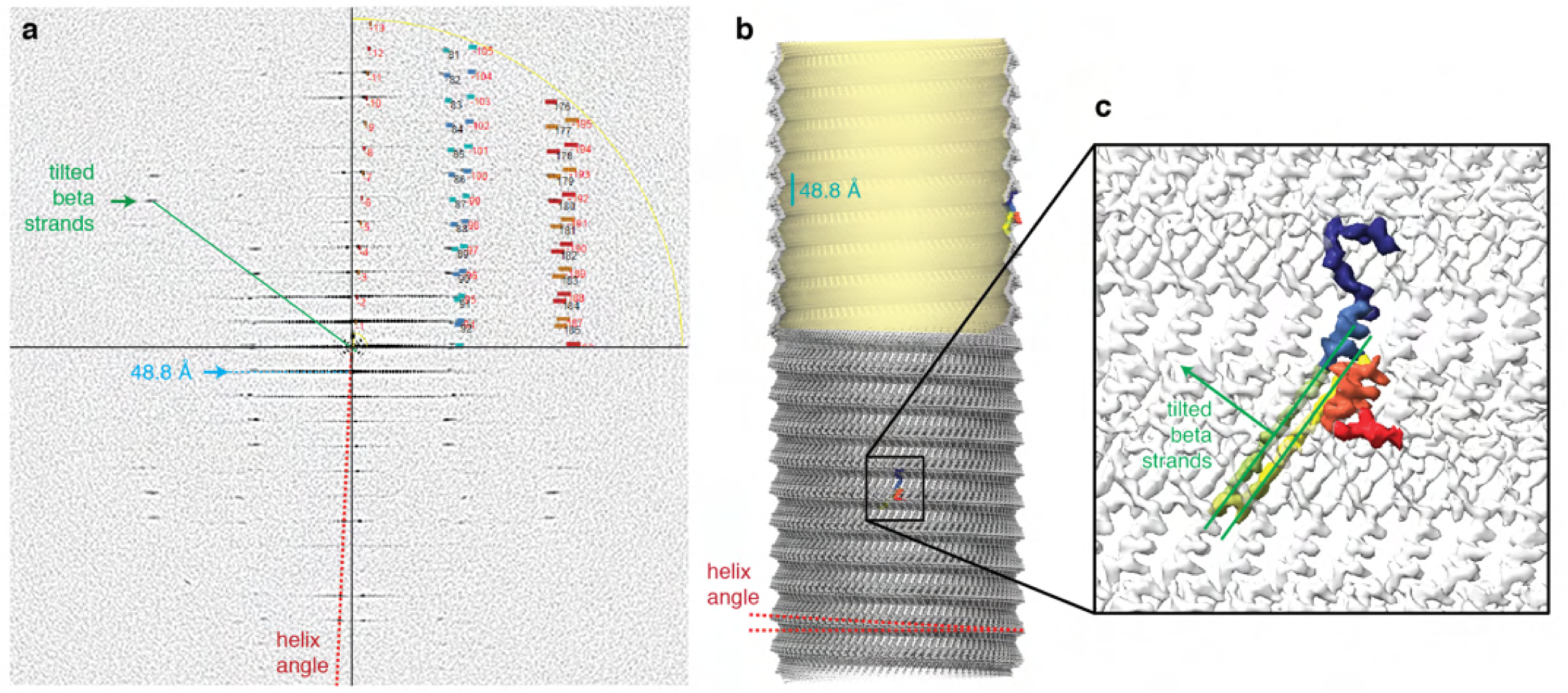
Summed in-plane rotated power spectra of final images used for cryo-EM reconstruction. (a) Power spectrum with first layer line at 48.8 Å, tilted β-sheet peak and helix angle annotated. The upper right quadrant is overlaid by a screenshot from Helixplorer (http://rico.ibs.fr/helixplorer/) used for interactive exploration of helical symmetry. Bessel function maxima correspond to final symmetry of 48.8 Å pitch and 92.93 units per helical turn turn. (b) cryo-EM density with pitch and helix angle annotated. (c) GvpA2 monomer with 36°tilted βstrands.

**Fig. S5.**
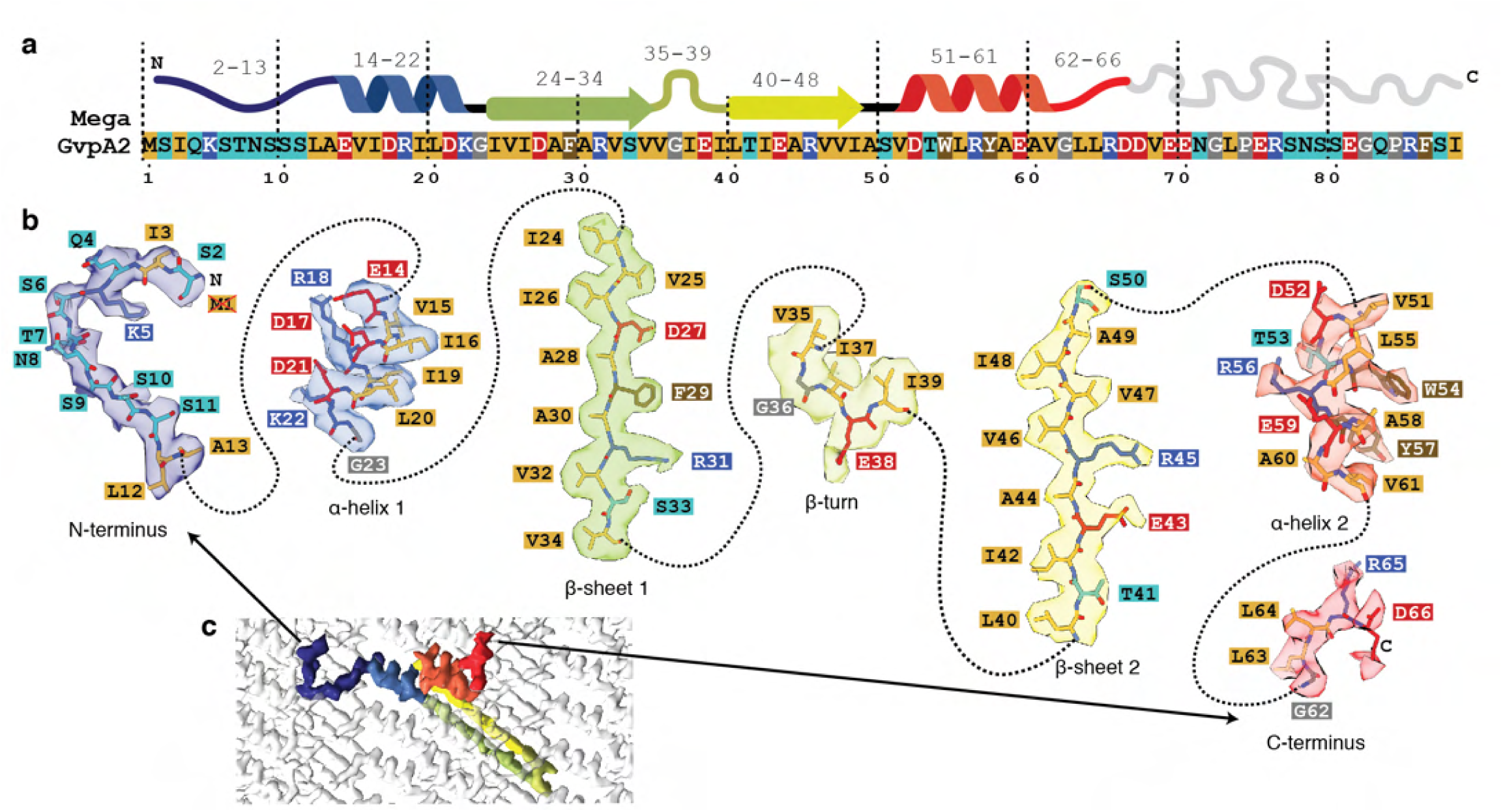
Details of the cryo-EM structure of the *B*.*megaterium* wall. (a) Primary and secondary structure of wall-forming protein GvpA2 (GvpB) from *B*.*megaterium* with physicochemical properties of side chains colour-coded (polar:turquoise, negative:red, positive:blue, hydrophobic:yellow) (b) Entire cryo-EM density of a monomer annotated with respective amino acid one-letter-codes and the atomic model. (c) Overview over cryo-EM density of a single monomer.

**Fig. S6.**
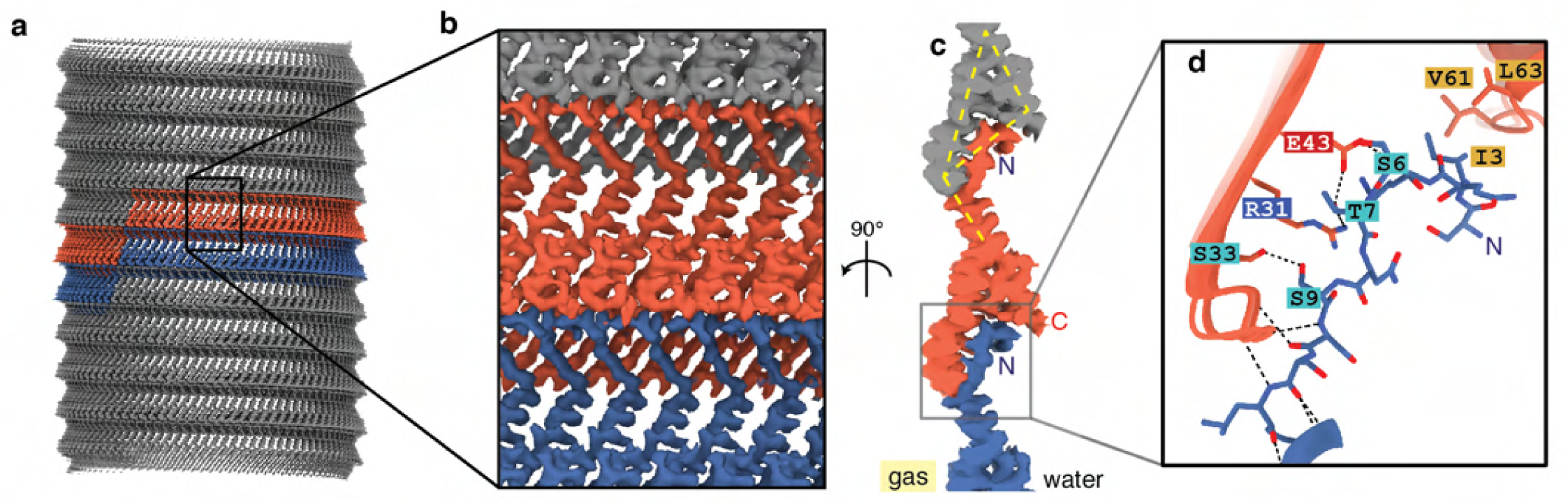
Details of rib-to-rib contact in the *B*.*megaterium* GV wall. (a) Cryo-EM density of the GV wall with two ribs highlighted in orange and blue. (b) Zoom-in to rib-to-rib contact (C) side-view (d) Molecular details of binding between N-terminus and the adjacent rib reveals hydrogen bonding between several residies as well as van-der-Waals interactions between I3 and V61,L63.

**Fig. S7.**
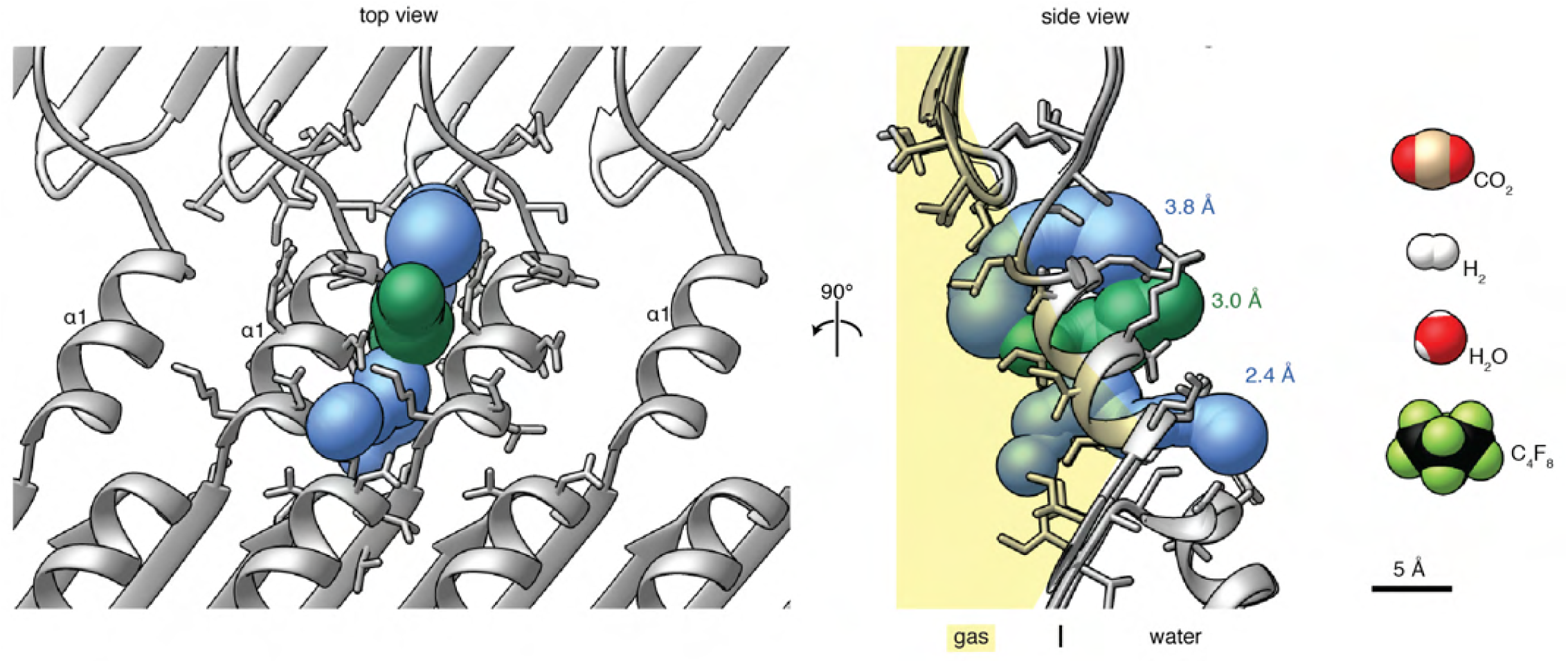
Gas pores in the *B*.*megaterium* wall. Top and side-view of the gas vesicle wall reveals slits between α-helices 1. The slit was modelled as adjacent tunnels (blue and green) in MOLE 2.5 (23) and shows gas-permeable openings of up to 3.8 Å diameter. The van-der-Waals surfaces of several gas molecules known to diffuse through the wall are shown to scale, with perfluorocyclobutane being the largest with 6.3 Å collision diameter (9).

**Fig. S8.**
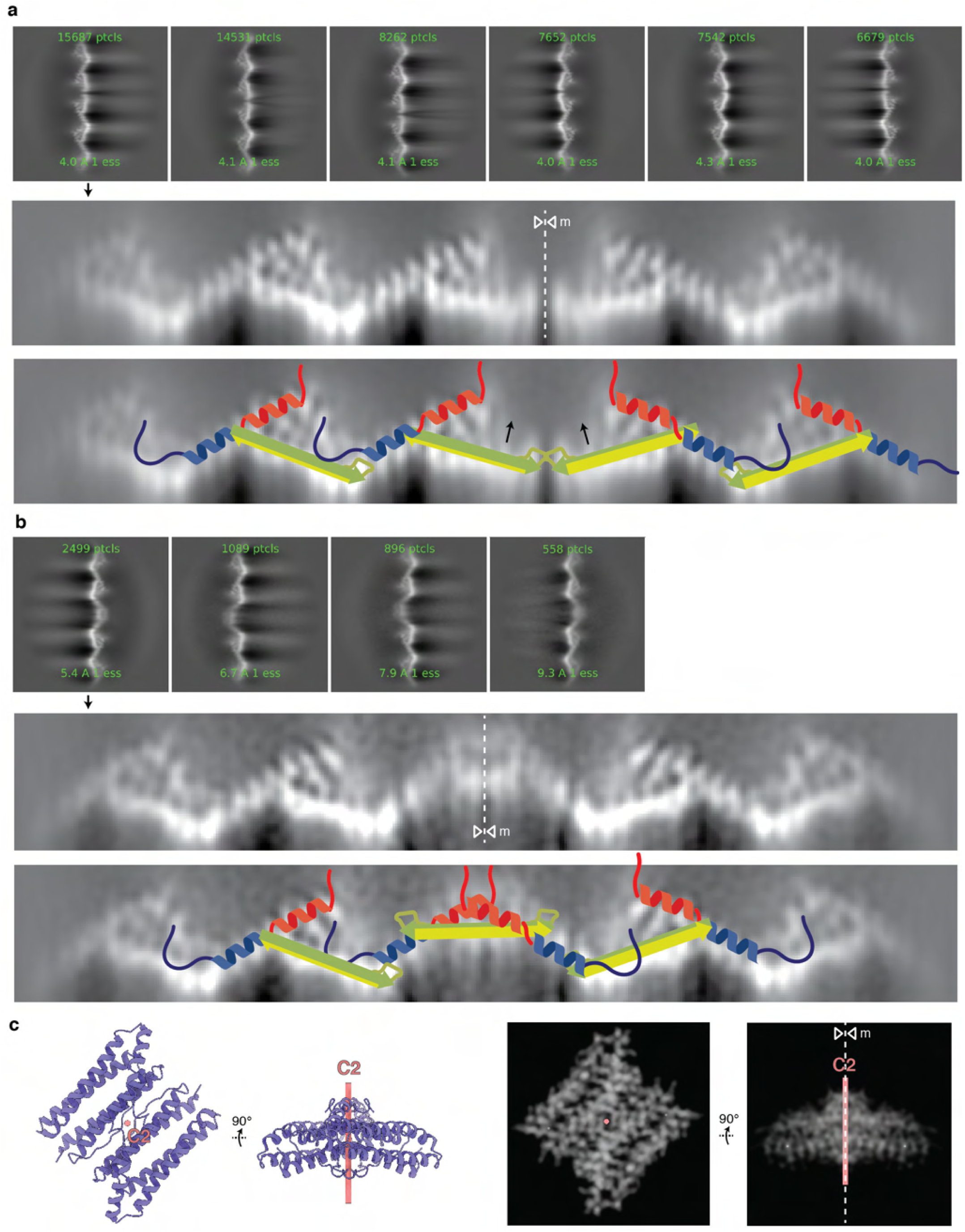
2D classification of *B*.*megaterium* seams. (a) 2D classes from the seam show perfect or near-perfect mirror symmetry. β-hairpins seem to hinge upwards at the seam (black arrows). (b) 2D classes from the putative polarity reversal point. Selected classes were magnified and sharpened for easier depiction. The mirror axis is shown (m, dotted white line). A cartoon is drawn on the 2D classes to visualise GvpA molecules with the N-terminus in blue and the C-terminus in red. (c) Demonstration of the fact that projection views orthogonal to a 180 degree rotation axis show mirror symmetry. Apoferritin dimer (pdb: 7ohf) is shown along the C2 axis and orthogonal to the C2 axis. 2D projection images (right) orthogonal to the C2 axis show mirror symmetry.

**Fig. S9.**
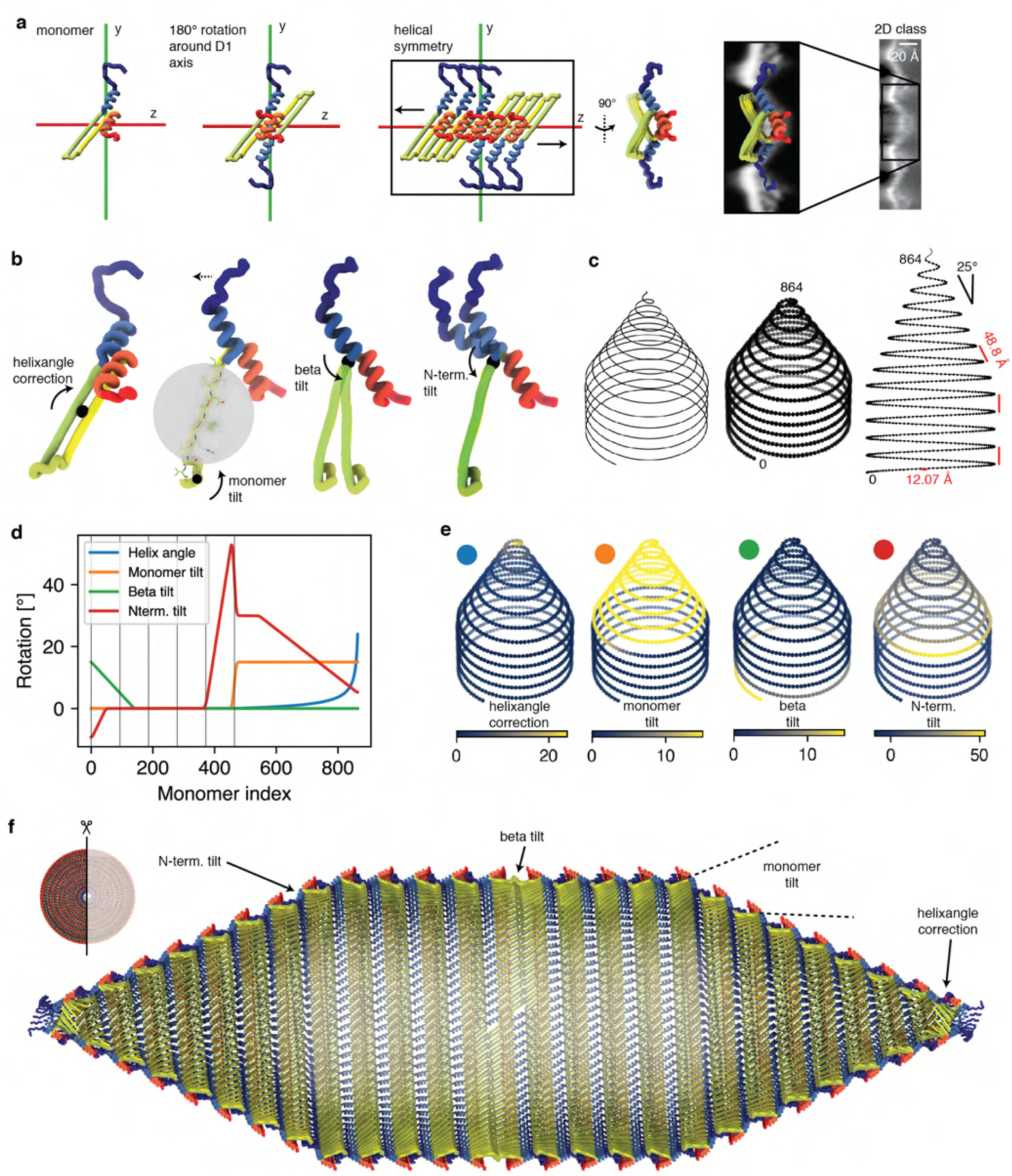
Construction of pseudo-atomic model of a whole *B*.*megaterium* GV. (a) A GvpA2 monomer was placed next to the x axis such that a 180 symmetry operation would reproduce a view corresponding to the experimental 2D class average. The β-sheets meet in an angle at this stage, which is later corrected by tilting the sheets. (b) Four rotation parameters (helixangle correction, monomer tilt, βsheet tilt and N-term. tilt) used in the model are visualised. (c) The model is based on a helical curve in space with a linearly decreasing radius in the cones. The pitch in both the cylinder and cone is 48.8 Å. 835 monomers are placed equidistantly on the curve with a distance of 12.07 Å. (d) Helixangle was extracted from the curve, while the other three parameters were manually tuned to fit the 2D class average data, make the βsheets line up in the polarity reversal point and avoid gaps by adjusting the N-termini. Vertical lines every 93 units depict the monomers on each wrung of the cylindrical part. (e) Parameters mapped onto the monomers (f) Cut-through of the final model highlighting the impact of the four model parameters.

**Fig. S10.**
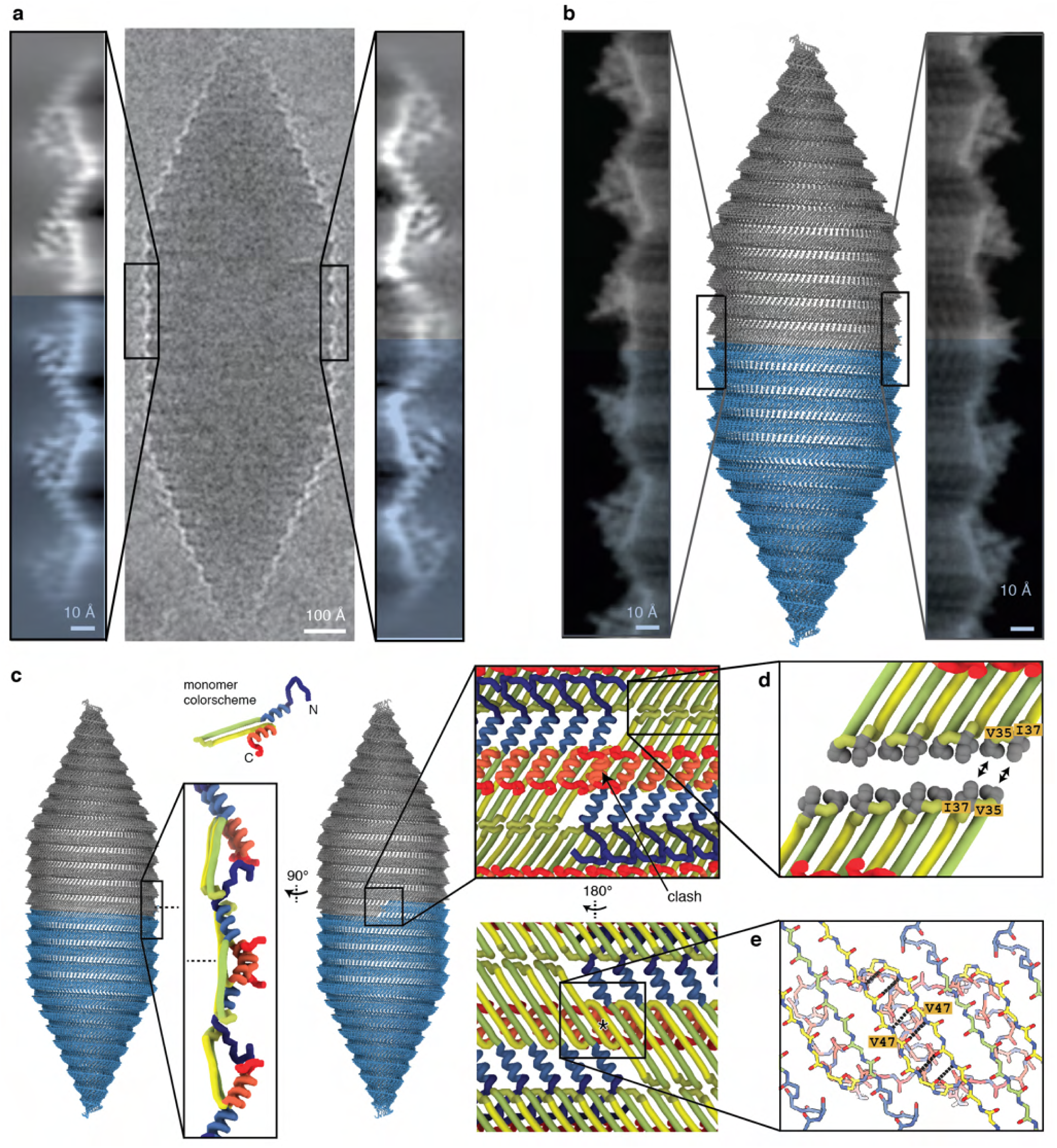
Pseudo-atomic model of *B*.*megaterium* gas vesicle with polarity reversal point. (a) 2D class from the seam and polarity reversal point and (b) simulated electron density from the pseudo-atomic model are in close agreement. (c) Detailled side-view and top-view of the polarity reversal point (PRP) of the GvpA rib with colorscheme indicating main-chain of the GvpA2 monomer. An unresolved clash between α2-helices from GvpA monomer next to the PRP is visualised. (d) In the model, the two oppositely rotated monomers next to the PRP are connected with 6 hydrogen bonds between the backbones of the β-sheets β2, with amino acid V47 being located around the D1-symmetry axis.

**Fig. S11.**
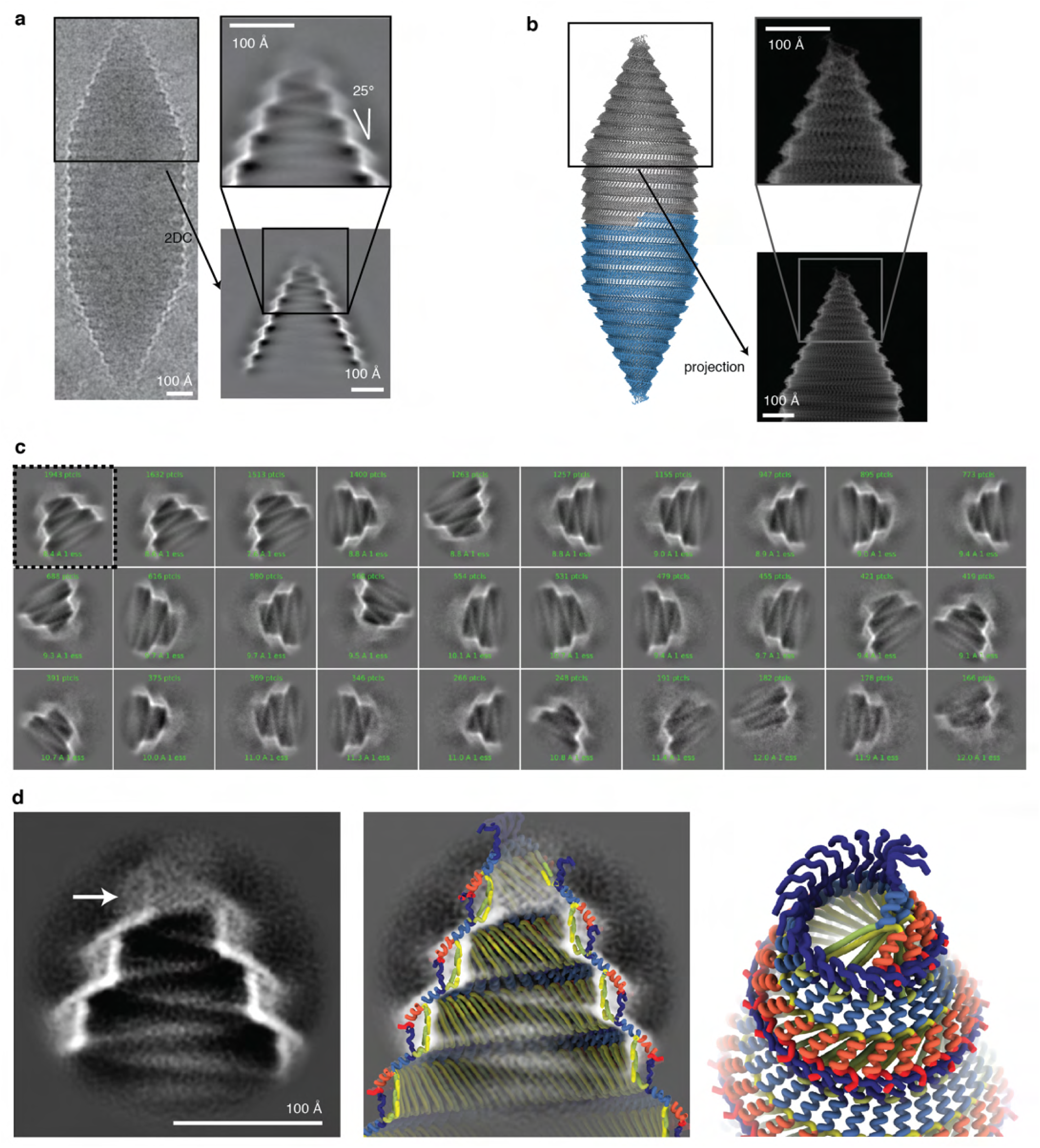
Structural analysis of gas vesicle tips. (a) Single cryo-EM image of a *B*.*megaterium* gas vesicle. The constrast is inverted to make electron density white. A 2D class average of gas vesicle tips reveals a linear decrease in radius at the tips with a cone angle of 25°. (b) Pseudo-atomic model of a GV with simulated 2D projections of the tip, closely matching the experimental data. (c) 2D classes of GV tips with smaller box size reveal more detail, but all end in a blurry density at the tip. Alignment of secondary structure features is not possible and indicated strong structural heterogeneity at the tips. (d) Zoom-in of one class average indicated in (c) (dotted line). Cut-through of pseudo-atomic GV model is fitted into the 2D class. The simplifed model of a helix with linearly decreasing radius breaks down at the very tip (arrow), leading to clashes between the main chains. N-termini from GvpA monomers at the tip come into close proximity and might close off the gas space.

**Fig. S12.**
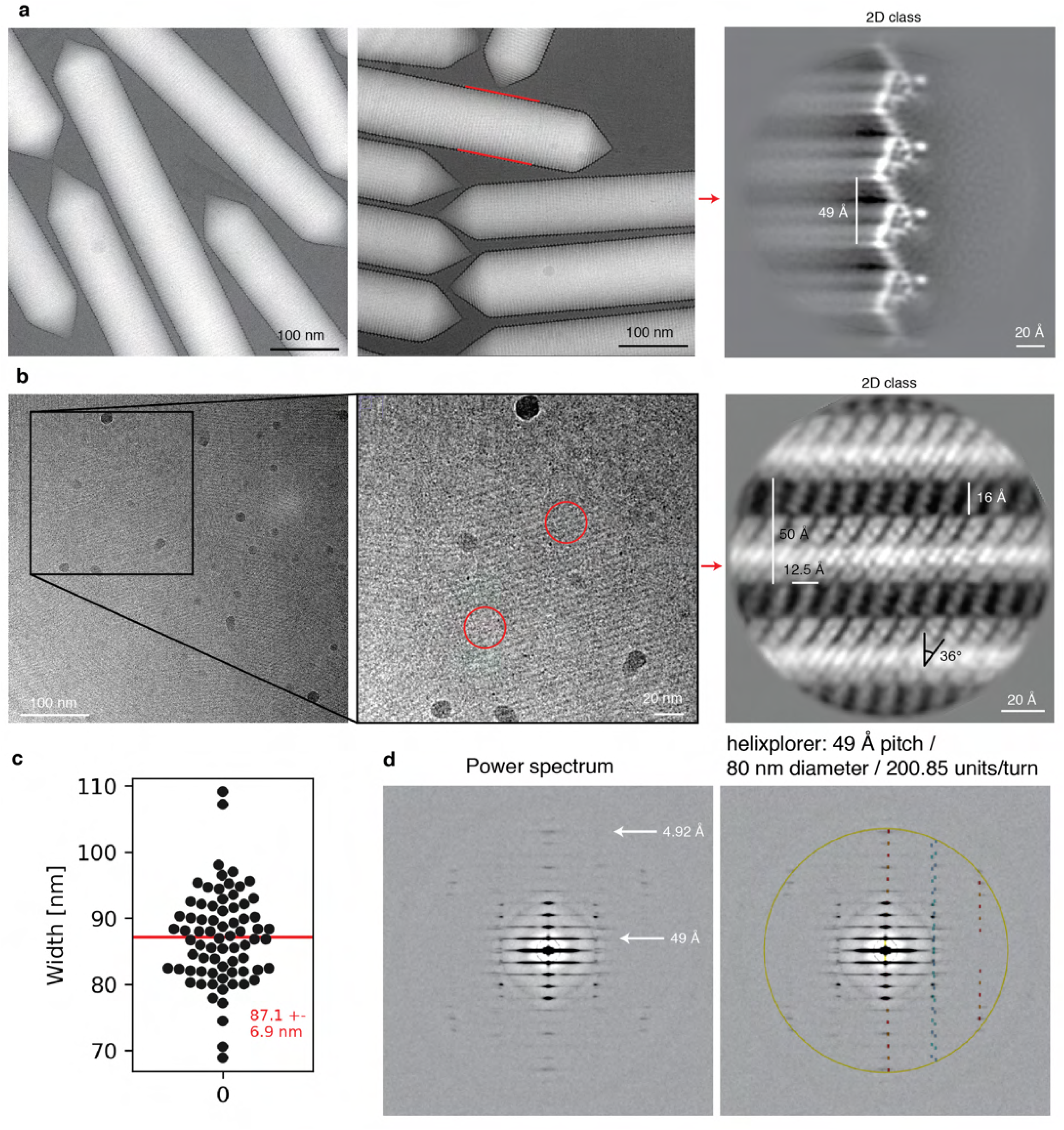
Cryo-EM of *A*.*flos-aquae* GVs. (a) Representative micrographs of *A*.*flos-aquae* gas vesicles. GV edges were analysed by 2D class averaging to give a low-noise highresolution 2D view of the edges, to reveal a repetetive zig-zag pattern. (b) The same dataset contained collapsed gas vesicle wall segments. Those can be averaged as well by 2D class averaging to reveal a high-resolution top-view of the GV wall. (c) Computing the sum of in-plane rotated power spectra of segments of all GVs in the dataset gives rise to a layer-line pattern typical for helical assemblies. This approach can be seen as a form of fiber diffraction where fibers are aligned computationally. Overlay with computed layer line patters (Helixplorer, http://rico.ibs.fr/helixplorer/) shows good agreement to a helix with 49Å pitch and 200.85 units per helical turn.

**Fig. S13.**
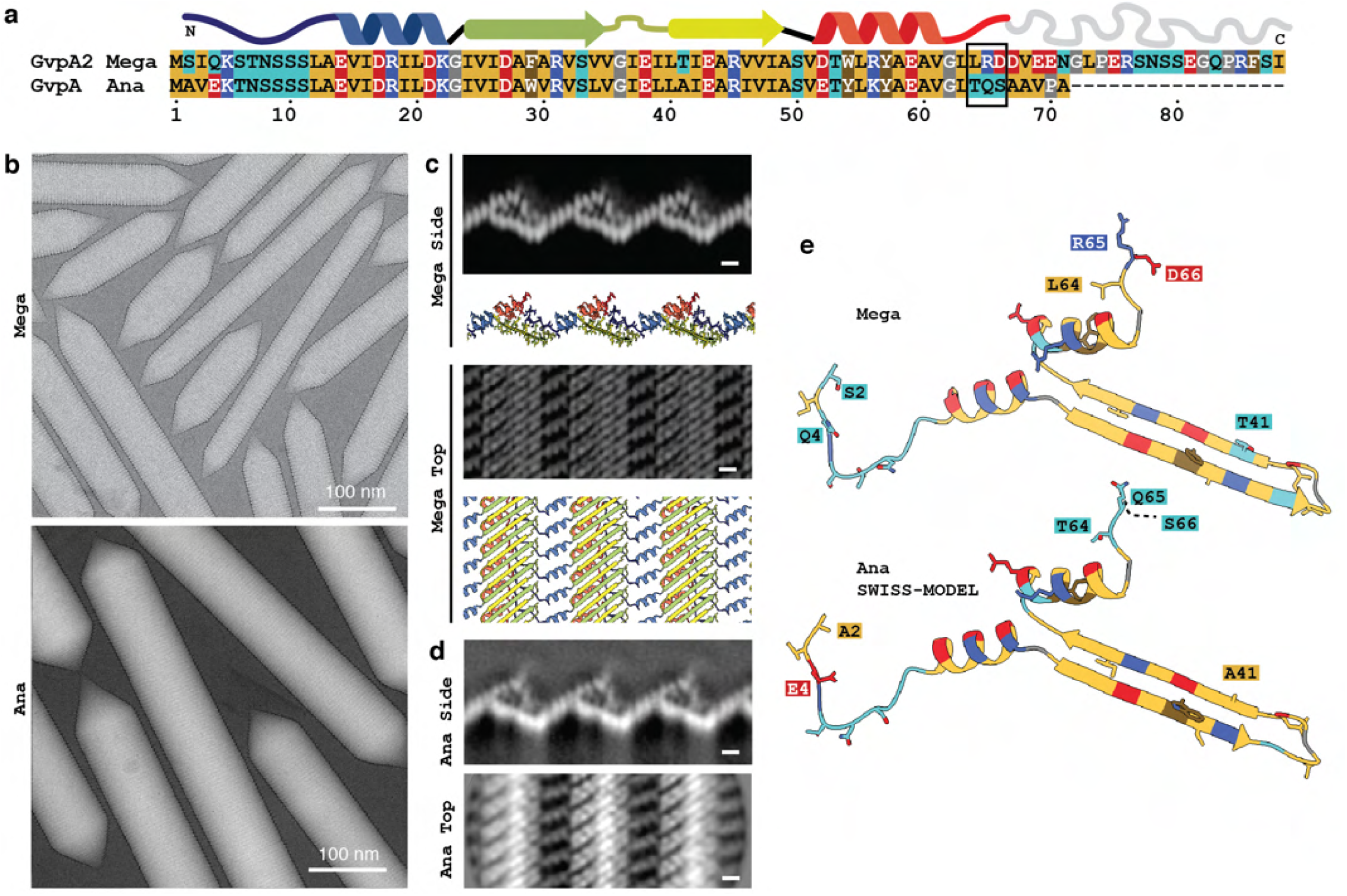
GvpA from *B*.*megaterium* and *A*.*flos-aquae* adopt the same fold and assembly. (a) Protein sequence of wall-forming protein GvpA from both *B*.*megaterium* and *A*.*flosaquae* are very similar (b) Representative cryo-EM micrographs of both gas vesicle types. (c) Projected side views and top views from the solved cryo-EM structure of *B*.*megaterium* GVs. (d) 2D classes from *A*.*flos-aquae* of the same views reveal very similar assembly structure. (e) Homology model of *A*.*flos-aquae* GvpA compared to *B*.*megaterium* with deviating residues displayed and property-changing mutations highlighted by one-letter-code. Residues are coloured according to side-chain chemistry as indicated in (a).

**Fig. S14.**
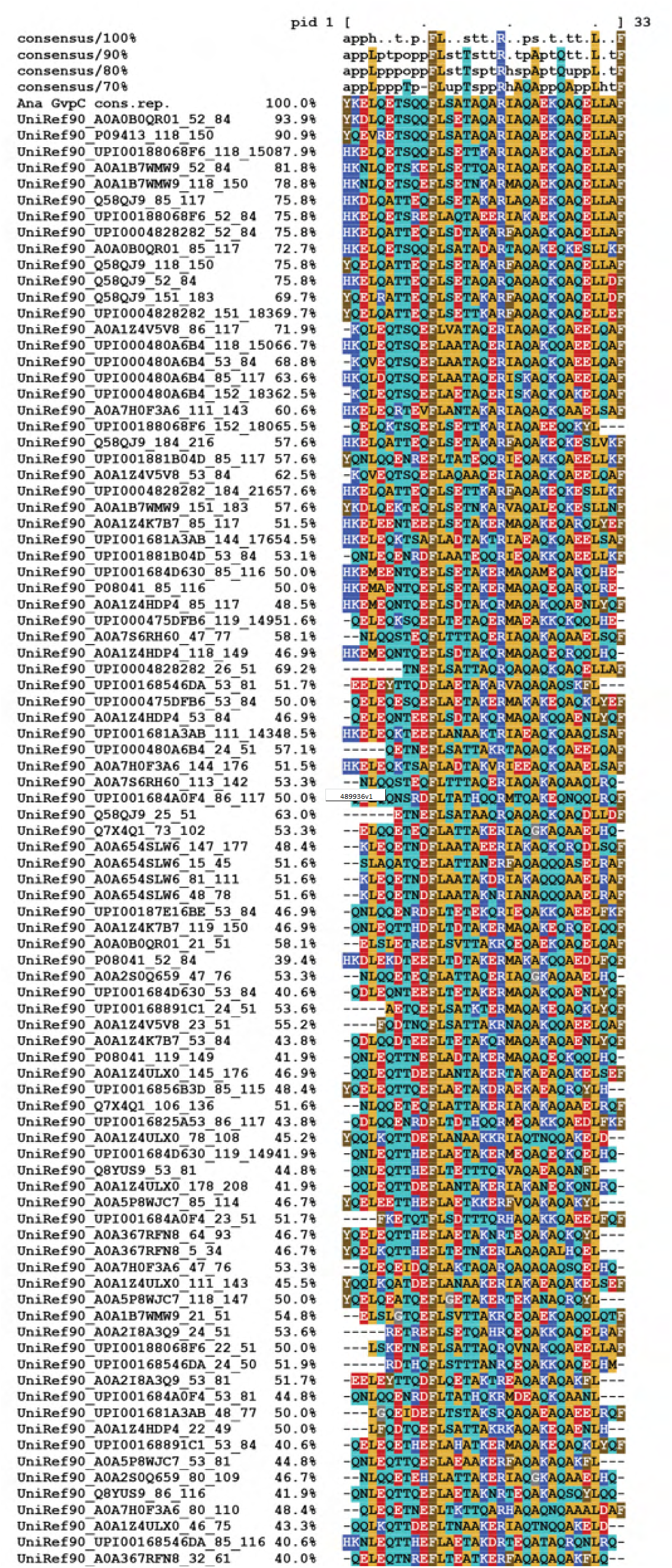
MSA of sequences similar to *A*.*flos-aquae* GvpC 33 AA repeats. Alignment of 33 amino acid repeat consensus sequence of Ana GvpC with 91 similar sequences from UniRef90 clusters with ∼40-100% sequence identity reveals a highly conserved pattern including leucine, phenylalanine and arginine residues. The consensus sequences on top were computed in MView and show residues conserved on at least 70 %, 80 %, 90 % and 100 % of sequences.

**Fig. S15.**
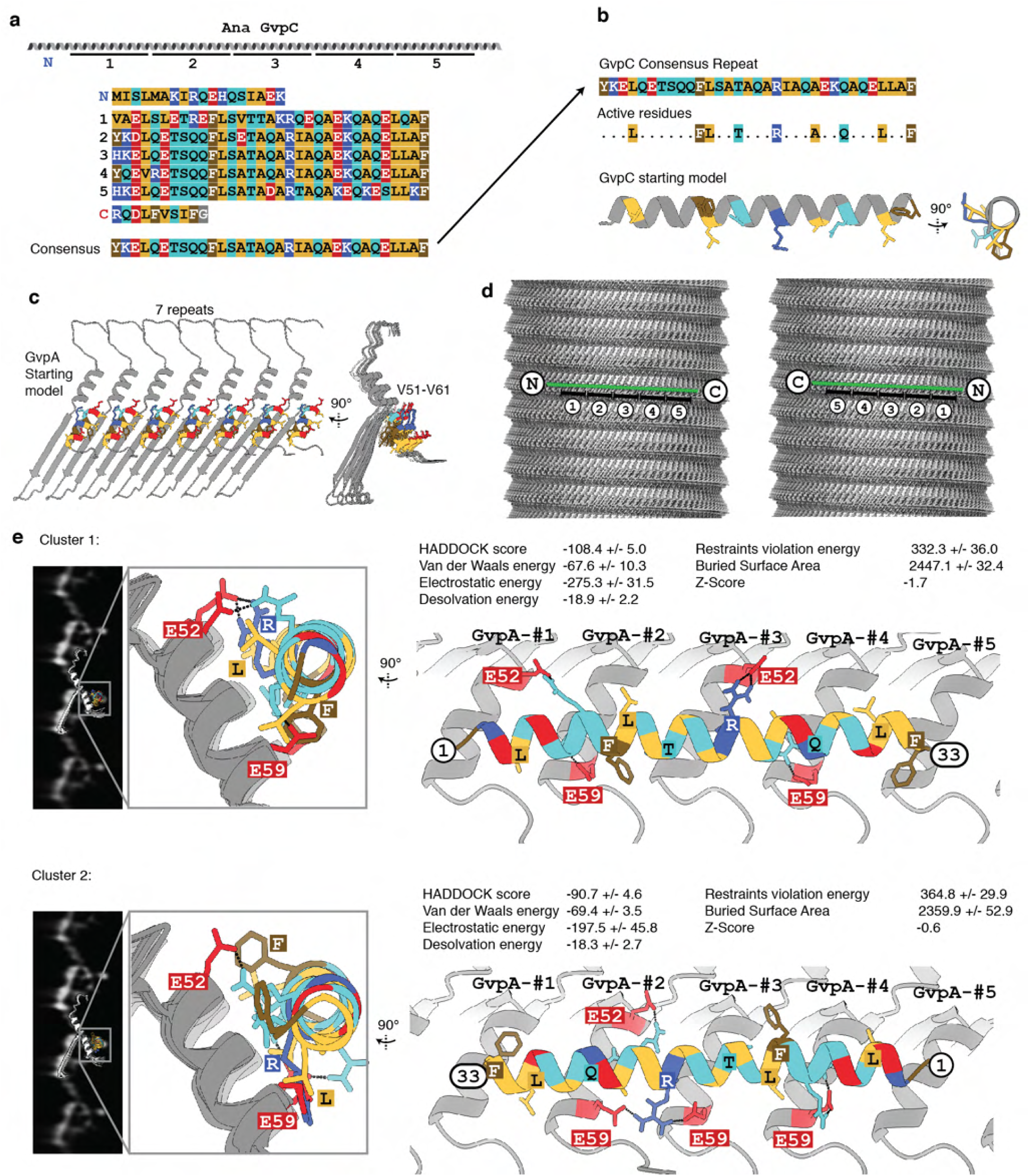
Binding analysis of GvpC to *A*.*flos-aquae* gas vesicles. (a) Primary sequence of Ana GvpC with 5 repeats and the consensus sequence of the repeats. (b) Identified highly conserved residues in the repeat are highlighted and chosen as ‘active residues’ in HADDOCK. A perfect α-helical peptide is used as the starting model. (c) Amino acids V51 to V61 on α-helix 2 are chosen as the active residues on GvpA based on observed close proximity in 2D class averages. GvpA was repeated 7 times according to the helical symmetry of the solved *B*.*megaterium* assembly. (d) Two possible binding geometries of GvpC are shown on a *B*.*megaterium* GV density oriented with the seam on the bottom and the tip on the top: direction from C to N-terminus is following the left-handed loop of the helix, or reverse (e) HADDOCK protein docking results between GvpC and GvpA fall into two main clusters, one highest-scoring solution with a score of -108.4 and a solution with reverse binding polarity of GvpC with a score of -90.7. The best solution of each cluster is shown. The model was fitted into 2D class averages of *A*.*flos-aquae* edges.

**Fig. S16.**
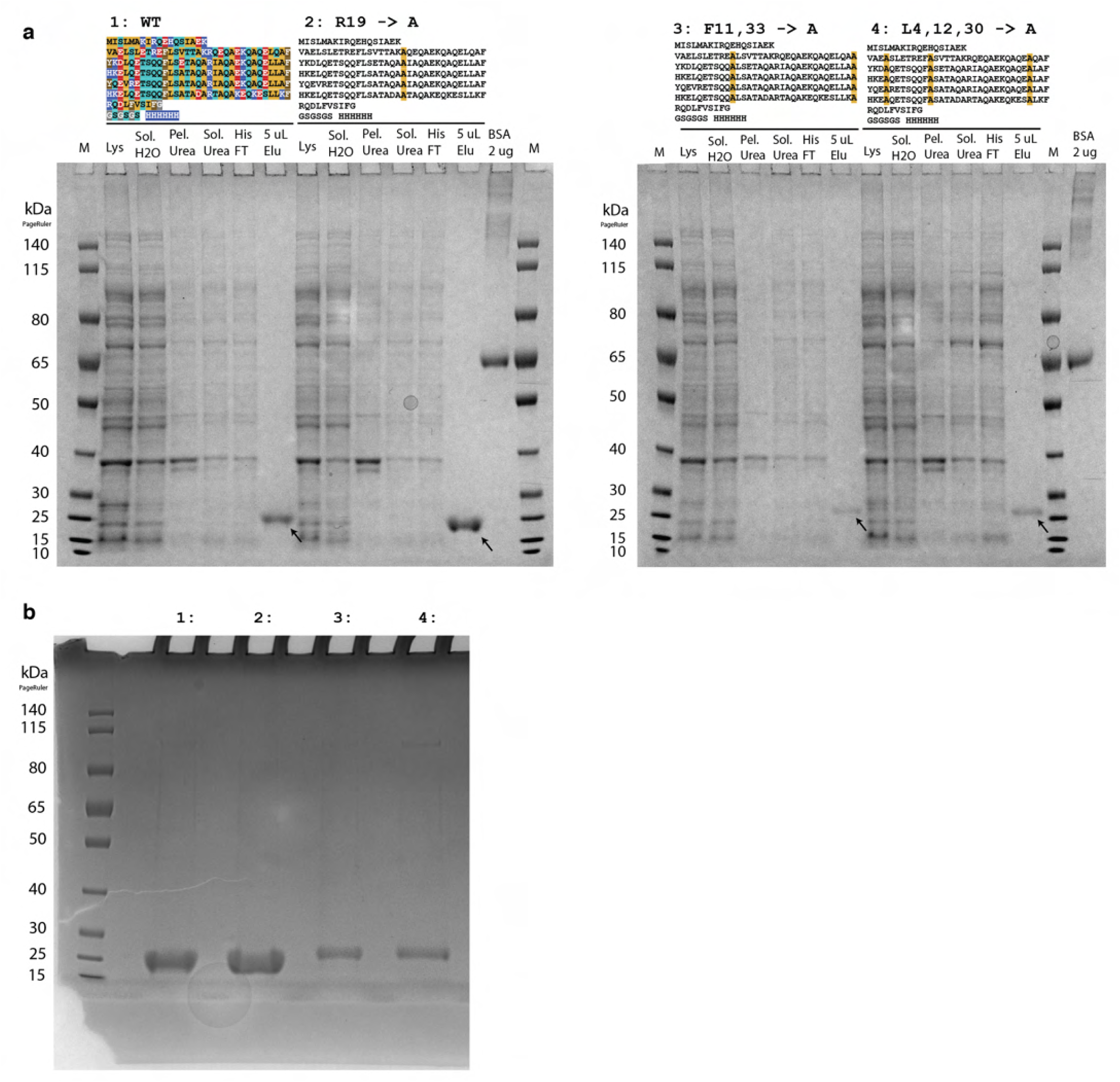
Purification of GvpC point mutants. (a) SDS-PAGE following purification steps of wild-type GvpC and point mutants. Primary sequence with mutation sites are indicated above the gel. Arrow indicates final product. Legend as follows: P:Pellet / SolH2O:Soluble fraction in aqueous solution / Pel.Urea:Pelleting fraction in urea-containing buffer / Sol.Urea:Soluble fraction in urea-containing buffer / HisFT:Flowthrough from Ni-NTA column / 5uLElu:Eluted fraction from Ni-NTA column (b) Overloading a SDS-PAGE gel with GvpC mutant sample indicates high degree of purity.

**Fig. S17.**
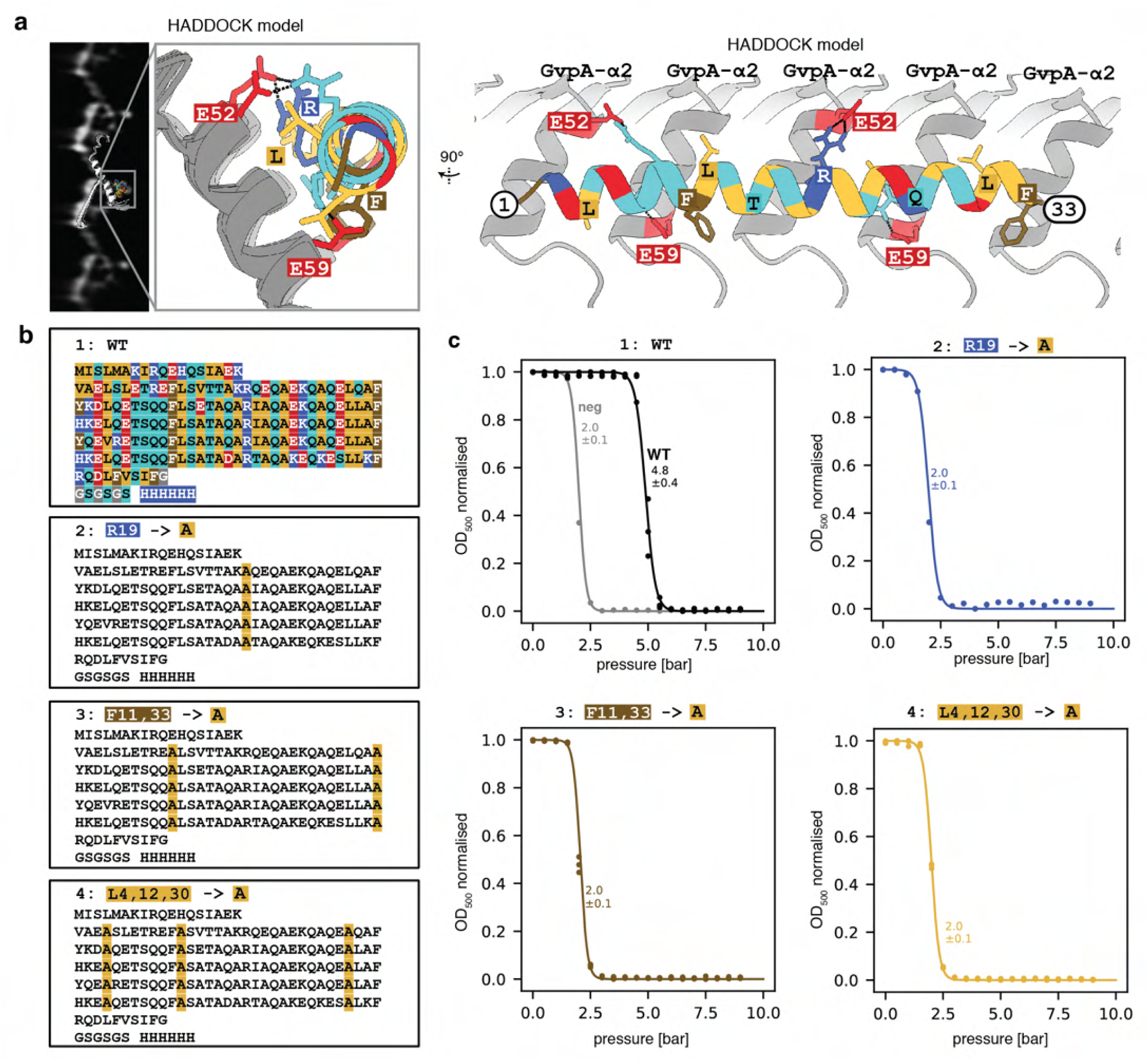
Mutation analysis of GvpC binding. (a) Best-scoring HADDOCK docking solution of GvpC binding to α-helix 2 of GvpA ribs. Highly conserved F, L and R residues are shown. (b) A wild-type construct and three mutants were designed with F, L and R residues in all five repeats mutated to alanine. All constructs have a C-terminal GSGSGS linker and 6x His-tag. (c) Collapse pressure measurements of the four constructs confirm binding to *A*.*flos-aquae* gas vesicles of the wild-type construct and loss of binding for the mutants. Stripped GVs before readdition of GvpC were measured as a negative control. Circles are measurement points from three independent readdition experiments. Solid curves are fits of a sigmoid function, with the stated number being the pressure when the normalised *OD*500 drops to 0.5.

## Notes

https://www.ebi.ac.uk/pdbe/entry/emdb/14238

https://www.ebi.ac.uk/pdbe/entry/emdb/14340

https://www.ebi.ac.uk/pdbe/entry/pdb/7r1c

https://doi.org/10.5281/zenodo.6458345

### Competing Interest Statement

The authors have declared no competing interest.

